# Bioengineering Developmentally Inspired Matrix Vesicles as Designer Nanotherapeutics for Bone Regeneration

**DOI:** 10.1101/2025.10.23.684111

**Authors:** Flurina Staubli, Paula Sobrevals Alcaraz, Robert M. van Es, Harmjan R. Vos, Eelco Bergsma, Debby Gawlitta, Kenny Man

## Abstract

Extracellular vesicles (EVs) are emerging as promising acellular nanotherapeutics for musculoskeletal repair. Matrix vesicles, a matrix-bound subset of EVs, are essential mediators of endochondral ossification in bone development and fracture repair. This study aims to design bioengineered matrix vesicles from hypertrophic cartilage microtissues to drive endochondral ossification for bone repair. Human mesenchymal stromal cell (hBMSC) microtissues were differentiated with/without BMP2 in chondrogenic or hypertrophic medium. Isolated matrix vesicles were characterized for physiochemical properties and biological functionality. BMP2 and hypertrophic conditioning significantly increased vesicle yield (1.5-fold), alkaline phosphatase activity (3.24-fold), calcium binding capacity (8.82-fold), and growth factor content (BMP2, VEGF). These vesicles promoted proliferation, migration, and mineralization of hBMSCs and enhanced angiogenesis in human endothelial colony forming cells (hECFCs), with BMP2 and hypertrophically conditioned vesicles showing the most pronounced effects. Proteomics analysis confirmed the enrichment of proteins involved in extracellular matrix remodelling, mineral deposition and vascularization within these hypertrophically engineered vesicles. These findings demonstrate that hypertrophic induction of cartilaginous microtissues substantially improves the yield and therapeutic potential of matrix vesicles. Taken together, this research unveils a powerful strategy to bioengineer developmentally inspired vesicles that not only recapitulate key cues of endochondral ossification but offers a tailorable, multifunctional nanotherapeutic platform for improved bone regeneration strategies.

**Graphical abstract:** 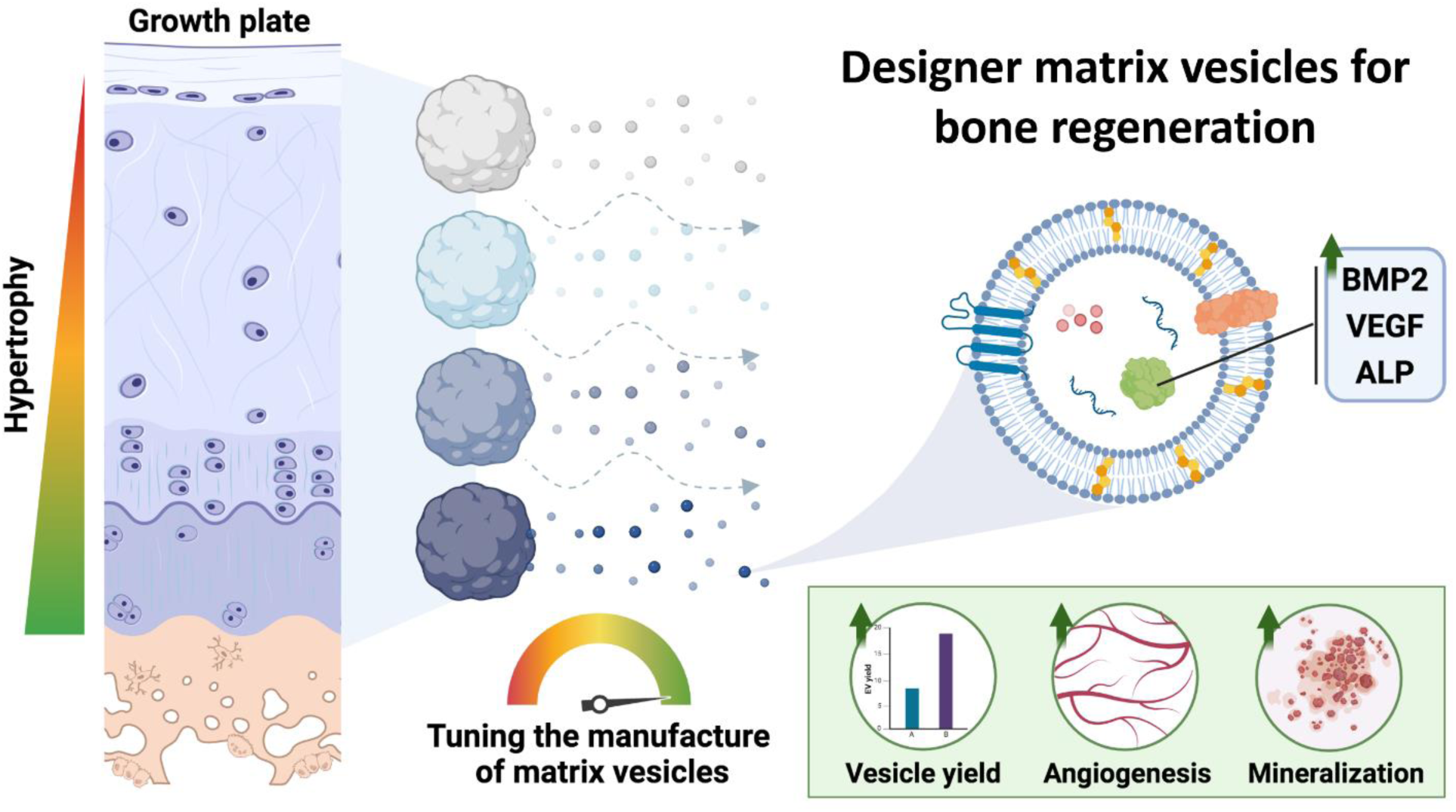

## 1. Introduction

Damage to bone can be caused by traumatic injury, tumor resection, congenital disorders, infection or age-related disorders such as osteoporosis (OP)[1]. Although bone tissue has a tremendous capacity to regenerate, in 10% of cases bone fractures cannot heal without surgical intervention[2]. The socioeconomic burden of these bone-associated disorders is expected to further increase due to our growing ageing population. To illustrate, annual fractures are projected to increase from 1.9 million to 3.2 million (68%), from 2018 to 2040, with related costs rising from $57 billion to over $95 billion[3]. The current gold standard treatment for bone defects, autografts, is associated with several challenges including morbidity and significant pain at the graft donor site[4, 5]. Moreover, it may not be possible to harvest sufficient amounts of autograft bone to reconstruct excessively large defects, commonly associated with cases of tumor resection or major trauma[6]. Patients that receive an alternative treatment, allogeneic (i.e. donor-derived) grafts, often experience complications including immune rejection, infection, fracture, and disease transmission[6, 7]. Biomaterials have emerged as a potential solution to overcome these limitations. These synthetic or processed natural materials can provide structural support and a scaffold for bone growth[8]. However, biomaterials alone often lack the biological cues necessary to stimulate optimal bone regeneration, limiting their effectiveness in complex bone repair scenarios. To enhance the functionality of biomaterials, researchers have explored incorporating osteoinductive growth factors, such as bone morphogenic protein 2 (BMP2), into bone graft substitutes. While this approach has shown positive clinical outcomes, it is not without risk. The supraphysiological concentrations of BMP2 required have previously resulted in severe complications such as heterotopic ossification, inflammation and myelopathy, which often require surgical intervention[9]. Furthermore, the use of BMPs following tumor resection is avoided due to the potential interaction of BMPs and cancer cells[10]. Thus, there is an urgent need for effective clinical solutions to repair damaged bone.

A well-established approach in bone tissue engineering emulates the process of fetal bone development and secondary fracture repair, which involves a cartilaginous intermediate stage and is known as endochondral bone formation[11]. This regenerative strategy focuses on implanting engineered cartilage tissues, which are subsequently remodelled into bone within the body[12]. These cartilaginous constructs can be generated using the patient’s own mesenchymal stromal cells (MSCs) cultivated in high-density pellet cultures. Numerous studies have confirmed that these cell-based cartilage implants stimulate endochondral bone regeneration (EBR) *in vivo*[13, 14]. Whilst these results are promising, the translation of cell-based therapies to the clinical arena are hindered due to several issues including regulatory hurdles, low cell survival, immune rejection, uncontrolled differentiation and teratoma formation[15, 16]. There is growing evidence that the beneficial effects exerted by cell therapies are partially caused by the paracrine factors they secrete, which have numerous trophic and immunomodulatory effects[17–19].

Of these factors, extracellular vesicles (EVs) are considered one of the most important secretory products, involved in numerous biological processes[20]. EVs are defined as cell-secreted lipid nanoparticles that contain a diverse biological cargo of nucleic acids, proteins and bioactive molecules[21]. The intercellular communication via the delivery of these EV-associated bioactive factors is critical in mediating biological functions between cells[22]. Excitingly, EV-based therapies possess numerous advantages compared to cell therapies, such as enhanced safety, improved physiochemical stability, targeted delivery, crossing biological barriers (transcytosis) and lower immunogenicity compared to their parent cells[21, 23, 24]. Additionally, vesicles exhibit improved clinical applicability when compared to their cell counterparts due to their non-living status and nanosize. When compared to similarly sized synthetic drug delivery systems, EVs exhibit improved circulation times, lower clearance rates and a decreased risk of systemic toxicity[25]. Moreover, the integration of EVs with biomaterials presents an exciting opportunity to create advanced functional materials for bone regeneration. EVs can be incorporated into various scaffold materials, such as hydrogels, nanofiber scaffolds, or 3D-printed composite structures. This combination allows for the controlled release of bioactive molecules contained within the EVs, while the scaffold provides structural support and potentially enhances the retention and efficacy of the EVs at the site of injury[8, 26].

Within the context of bone, EVs play crucial roles in various physiological processes. Growing evidence suggests that EVs are intrinsically involved in regulating bone homeostasis by mediating intercellular communication. These cell-derived nanoparticles are also fundamentally important in bone development, as matrix-bound vesicles are critical for endochondral ossification (EO) [23, 27]. Furthermore, EVs contribute significantly to bone repair processes. During the repair of mechanically stable fractures, hypertrophic cartilage within the fracture callus undergoes calcification, vascularization, and remodelling into woven bone. In this context, matrix vesicles (MtVs), a specific type of membrane-bound EVs, are secreted by hypertrophic chondrocytes and function to nucleate cartilage calcification and stimulate vascular ingrowth[28]. In more detail, these MtVs consist of lipids and proteins which chelate Ca^2+^ and P_i_, facilitating mineralization[29, 30], in addition to harbouring angiogenic bioactive molecules such as vascular endothelial growth factor (VEGF)[31]. Moreover, growing evidence suggest that MtVs are phenotypically distinct to media secreted vesicles due to their high tissue-nonspecific alkaline phosphatase (TNAP) activity[32]. Thus, harnessing MtVs for regenerative medicine is an attractive nanoscale therapeutic approach to stimulate EBR. Several studies have reported the successful isolation of MtVs procured from animal-derived cartilage tissues[31, 33]. However, the clinical use of these nanoparticles is hindered due to issues with xenogeneic transplantation, such as severe immune responses and potential disease transmission[34]. While MtVs from human-derived cartilage tissues could be an alternative, their use is associated with drawbacks such as limited tissue availability, challenging scaleup and potential allogeneic immune rejection[35]. In contrast, cartilage tissues derived from chondrogenically differentiated MSCs offer significant advantages. MSCs can be expanded in culture, providing a more abundant and scalable source of tissue for EV production. Moreover, MSC-derived tissues are less likely to trigger immune rejection compared to native cartilage, even in allogeneic settings. This reduced immunogenicity is attributed to the immunomodulatory properties of MSCs, which are partially retained in their differentiated state and their secreted vesicles. Thus, by engineering MSC-derived cartilage tissues that closely mimic the hypertrophic extracellular matrix (ECM) composition and structure of the soft callus, we may enable the scalable production of soft callus-mimetic MtVs for EBR.

To the best of our knowledge, this is the first study to bioengineer soft callus-mimetic MtVs through refining the *in vitro* manufacture of MSC-derived hypertrophic cartilage microtissues. This bioengineering approach allows us to precisely control and tune the properties of the resulting MtVs, enabling the creation of customized EVs that closely mimic the characteristics of those found in natural soft callus. By refining this process, we aim to develop MtVs with tailored functionalities for specific applications in bone regeneration and repair. Our research contributes to the future development of advanced materials for bone regeneration. The engineered soft callus-mimetic MtVs show promise for biomaterial functionalization, potentially leading to improved scaffolds with capabilities beyond structural support. By incorporating these multifunctional MtVs, biomaterials could promote bone formation through the release of bioactive factors, interact with the local microenvironment, support vascularization, and facilitate endochondral bone regeneration. This approach combines nanotechnology and biomimetic design, offering new possibilities for bone regeneration strategies.

## 2. Materials and Methods

### 2.1. Study Design and Overview

Figure 1 illustrates the overall experimental outline of the manufacture of hypertrophic cartilage tissues and the MtVs’ characterisation/biological assessment. For the generation of cartilage microtissues, hBMSCs were differentiated in chondrogenic medium (+TGFβ, +/- BMP2) for 21 days. Then, microtissues were either maintained in chondrogenic medium (+TGFβ, +/- BMP2) or switched to hypertrophic medium for a further 7 days. After 28 days, the microtissues were characterized for the presence of chondrogenic/hypertrophic markers via histological staining and biochemical analysis. The obtained MtVs were characterized by nanoparticle tracking analysis, transmission electron microscopy (TEM), protein quantification, alkaline phosphatase (ALP) activity, CD63-, BMP2- and VEGF-ELISA, and proteomics analysis. The biological functionality of MtVs on hBMSC uptake, proliferation, osteogenic differentiation and human endothelial colony forming cell (hECFC) angiogenesis was evaluated.

**Figure 1.**
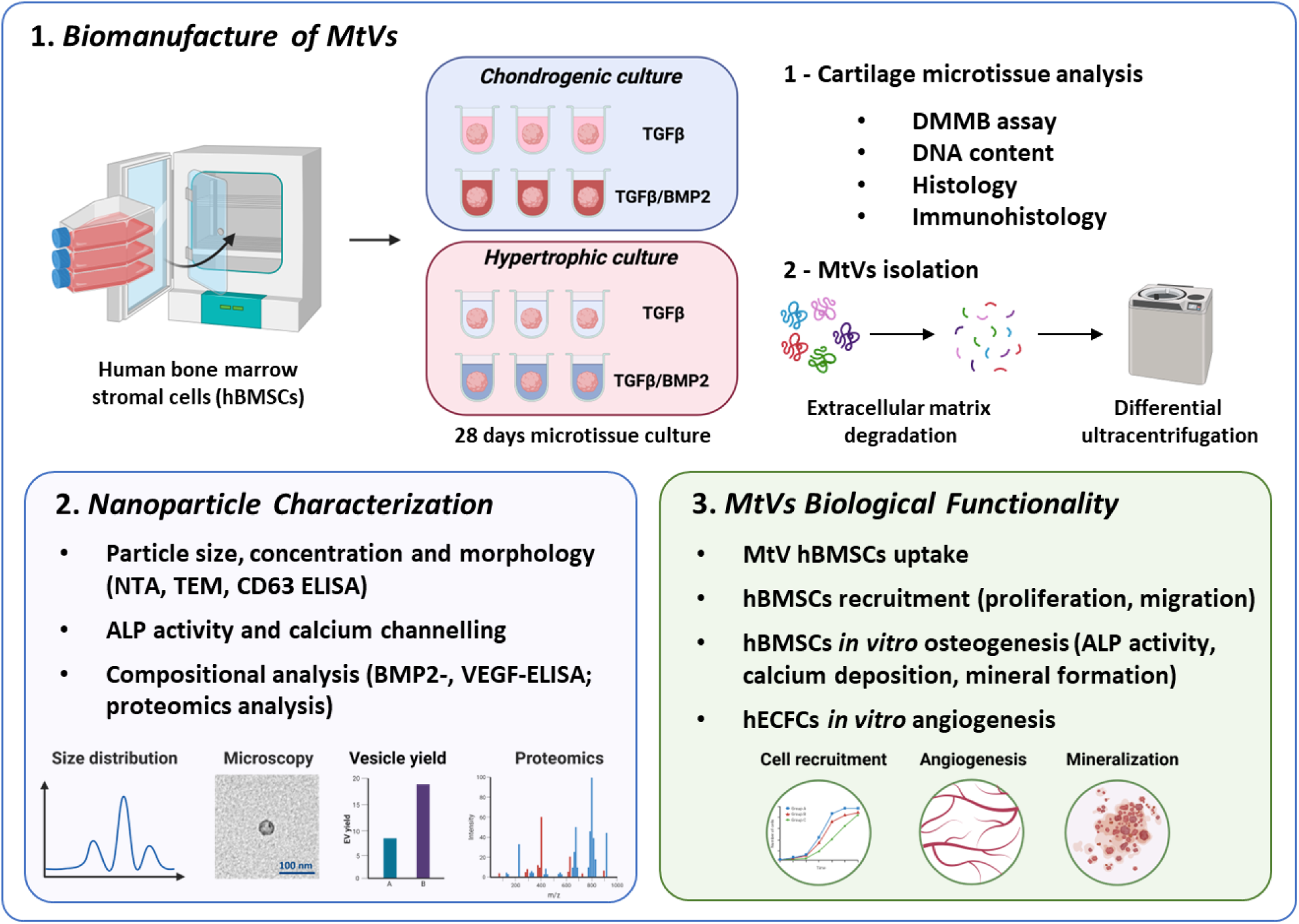
- Schematic representation of the experimental overview. hBMSC microtissues were generated and cultured in chondrogenic differentiation medium (+TGFβ, +/- BMP2) for 21 days. Following which, tissues were either maintained in chondrogenic medium (+TGFβ, +/- BMP2) or switched to hypertrophic medium for a further 7 days. After 28 days of culture, the presence of chondrogenic and hypertrophic markers was assessed via biochemical and histological analyses. Moreover, the cartilage microtissues were enzymatically digested and subjected to differential ultracentrifugation to obtain the MtVs. The size, morphology, quantity, ALP activity, BMP and VEGF content of these vesicles were assessed. The biological function of obtained MtVs on hBMSC uptake, proliferation, osteogenic differentiation and hECFC angiogenesis was evaluated *in vitro*.

### 2.2. Isolation and Expansion of hBMSCs

hBMSCs were isolated from bone marrow aspirates of patients (N=3; 15, 20, 20 years of age) after informed consent, in accordance with a biobank protocol approved by the local Medical Ethics Committee (TCBio-08-001, University Medical Center Utrecht)[36]. Adherent cells were maintained at 37°C under humidified conditions and 5% carbon dioxide (CO_2_) in basal medium consisting of α-MEM (22 561, Invitrogen) supplemented with 10% heat-inactivated fetal bovine serum (FBS, S14068S1810, Biowest), 0.2 mM L-ascorbic acid 2-phosphate (A8960, Sigma-Aldrich), 100 U/mL penicillin with 100 mg/mL streptomycin (15 140, Invitrogen), and 1 ng/mL basic fibroblast growth factor (233-FB; R&D Systems). The media were refreshed every 2 days until the cells reached 80% confluence. All experiments described in this study using hBMSCs were conducted at passage 4. For EV dosing experiments, EVs were removed from the FBS via ultracentrifugation at 120,000 g for 16 h prior to use in culture medium.

### 2.3 Generation of hBMSC-Derived Callus-Mimetic Microtissues

hBMSC microtissues were created as previously described[37]. Briefly, 250,000 hBMSCs in 200 µl of chondrogenic differentiation medium were transferred into each well of a 96 well round (U) bottom suspension plate and centrifuged at 300 g for 4 min. The chondrogenic differentiation medium consisted of high glucose Dulbecco’s modified eagle medium (DMEM, 31966, Thermo Fisher Scientific), 1% ITS (insulin-transferrin-selenium)+ premix (354352, Corning), 100 nM dexamethasone (D8893, Sigma-Aldrich), 0.2 mM L-ascorbic acid 2-phosphate, 100 U/mL penicillin with 100 mg/mL streptomycin, 10 ng/mL transforming growth factor-β1 (TGFβ1, Peprotech) with or without the addition of 100 ng/mL BMP2 (EU/1/09/526/001, InductOS). For the first 4 days, the medium was changed daily and afterwards every 3rd day. After day 21 of chondrogenic differentiation, half of the microtissues was cultured in hypertrophic medium, which consisted of DMEM (31966, Thermo Fisher Scientific), 1% ITS + premix (354352, Corning), 1 nM dexamethasone (D8893, Sigma-Aldrich), 0.2 mM L-ascorbic acid 2-phosphate, 100 U/mL penicillin with 100 mg/mL streptomycin, and 1 nM 3,3′,5-triiodo-L-thyronine (T2877, Sigma-Aldrich). Differentiation in chondrogenic or hypertrophic medium proceeded for 7 additional days till day 28. 4 microtissue groups were obtained: CT (chondrogenic medium/-BMP2), CB (chondrogenic medium/+BMP2), HT (hypertrophic medium/-BMP2), HB (hypertrophic medium/+BMP2).

### 2.4. Histological Analysis of hBMSC-derived Callus-Mimetic Microtissues

hBMSCs microtissues were washed with phosphate buffered saline (PBS) and fixed in 4% formaldehyde overnight at 4°C. Samples were then dehydrated in a series of increasing ethanol solutions (70 - 100%), followed by xylene and embedded in paraffin wax. Samples were sectioned (5 µm) using a microtome (Microm HM340E; Thermo Fischer Scientific). Safranin O and Toluidine blue staining was conducted to visualize glycosaminoglycans (GAGs) content. For Safranin O staining, the sections were deparaffinized, washed in deionized water and stained with Weigert’s hematoxylin. After samples were washed in running tap water and rinsed in deionized water, the sections were stained with fast green (0.4% w/v, Sigma-Aldrich) to visualize collagenous matrix (green). Sections were briefly washed in acetic acid (1% v/v), then incubated with Safranin O solution (0.125% w/v) to visualize proteoglycans (red). For Toluidine blue staining, sections were stained with Toluidine blue solution (0.4%, Sigma-Aldrich) in 0.1 M sodium acetate (pH 4, Sigma-Aldrich) to stain proteoglycans (purple) and then counterstained with fast green solution (0.2% w/v, Sigma-Aldrich). To assess calcium production, sections were incubated with 40 mM Alizarin Red S solution (pH 4.1, Sigma Aldrich) for 10 min. The unbound staining solution was removed by distilled water washes. To detect mineralization, von Kossa staining was performed by incubating samples in 1% silver nitrate (Sigma-Aldrich) for 1 h under a light bulb, followed by washing in 5% sodium thiosulfate (Alfa Aesar) and counterstained with haematoxylin. The stained sections were dehydrated in increasing ethanol solutions and incubated in xylene prior to mounting in Eukitt Quick hardening mounting medium (Sigma-Aldrich).

Immunohistochemical staining was conducted to visualize collagen type I, collagen type II, collagen type X, VEGF and ALP protein expression. Following the deparaffinization, sections were incubated with 0.3% v/v H_2_O_2_ to block endogenous peroxidase activity. For collagen type 1, collagen type II and ALP protein, antigen retrieval was conducted by incubation with pronase (1 mg/ml), followed by hyaluronidase (1 mg/ml) at 37°C for 30 mins each. For VEGF staining, antigen retrieval was carried out by incubating sections in 10 mM citrate buffer (pH 6) at 95°C for 20 mins. For collagen type X, antigen retrieval was performed by incubating sections in pepsin (1 mg/ml in 0.5 M acetic acid pH 2) at 37°C for 2 h, followed by hyaluronidase (10 mg/ml) at 37°C for 30 mins. Following antigen retrieval, samples were incubated with 5% bovine serum albumin-PBS at room temperature for 30 mins to block aspecific protein binding. The primary antibodies for collagen type II (0.6 µg/mL, II-II6B3), VEGF (5 µg/mL VG-1, ab1316), collagen type I (100 µg/ml, CP17-100, Omnilabo), collagen type X (10 µg/mL, X53; Thermo Fisher Scientific) and ALP (2 µg/mL, ab126820) were incubated with the sections at 4°C overnight. The secondary antibody, BrightVision HRP-anti-mouse IgG (VWRKDPVM110HRP), was applied followed by 3,3′-diaminobenzidine oxidation. Finally, samples were counterstained with hematoxylin prior to dehydration and mounting using the Eukitt Quick-hardening mounting medium. The stained sections were visualized using an Olympus BX51 microscope (Olympus DP73 camera, Olympus).

### 2.5. Biochemical Analysis

Samples for GAG, DNA and protein content analysis were digested at 60°C overnight in papain digestion buffer (250 µg/mL papain, 0.2 M NaH_2_PO_4_, 0.1 M EDTA and 0.01 M DL-cysteine hydrochloride; all from Sigma-Aldrich). Total GAG quantification was assessed using the 1,9-dimethyl-methylene blue (DMMB pH 3.0; Sigma-Aldrich) assay. Absorbance values were detected at 525 and 595 nm. DNA content was determined using the Quant-iT Picogreen dsDNA assay (P11496, Thermo Fisher Scientific) according to the manufacturer’s instructions. Protein content was quantified using the MicroBCA Protein Assay kit (Thermo Scientific) according to the manufacturer’s instructions.

### 2.6. EV isolation and Characterization

#### 2.6.1. Isolation of MtVs

Microtissues were incubated in collagenase II (500 U/ml; Sigma-Aldrich) for 3 h at 37°C. Digested microtissues were passed through a cell strainer (40 μm) and centrifuged at 2000 g for 20 min to remove cell debris. Leveraging size-dependent sedimentation, differential ultracentrifugation was used to isolate microtissue-derived MtVs, effectively procuring two distinct EV subpopulations based on size. It is common that a single population of EVs are isolated from cells/tissues of interest. In this study, we obtained both MtV subsets enabling a more thorough evaluation of the influence of BMP2/hypertrophic conditioning on the entire microtissue-MtVs population. EV supernatant was centrifuged at 10,000 g for 30 mins to obtain the large matrix vesicles (L-MtVs) subpopulation, whilst the supernatant was spun at 120,000 g for 70 mins. The EV pellet was washed with fresh PBS and spun for 120,000 g for 70 mins. The remaining EV pellet was resuspended in fresh PBS to collect the small matrix vesicles (S-MtVs) subpopulation. MtVs were stored at - 80°C until required. For both the L-MtV and S-MtV population, 4 EV groups were obtained: CT (chondrogenic medium/-BMP2), CB (chondrogenic medium/+BMP2), HT (hypertrophic medium/-BMP2), HB (hypertrophic medium/+BMP2). All ultracentrifugation steps were performed at 4°C using the Optima XE-90 Ultracentrifuge (Beckman Coulter) and a SW 32 Ti Rotor Assembly (Beckman Coulter). Microtissue conditioned media were collected at days 21 and 28 and EVs within the microtissue secretome (sEVs) were obtained following the ultracentrifugation steps described above.

#### 2.6.2. Particle Size and Concentration

Protein concentration of obtained MtVs, sEV and conditioned medium was quantified using the MicroBCA Protein Assay kit (Thermo Scientific) according to the manufacturer’s instructions. Nanoparticle tracking analysis (NTA) was conducted to evaluate particle size distribution and concentration using the Nanosight NS500 (NanoSight NTA 3.1). The CD63 concentration of MtVs was evaluated using the Human CD63 ELISA kit (RayBiotech) following the manufacturer’s instructions.

#### 2.6.3. Transmission Electron Microscopy (TEM)

EV imaging was conducted using the TFS Tecnai 20 transmission electron microscope. Samples were physisorbed to 200 mesh, carbon-coated copper formvar grids (Agar Scientific) and negatively stained with 1% uranyl acetate.

#### 2.6.4. MtVs ALP Activity

ALP activity of MtVs was evaluated using the p-nitrophenyl phosphate (pNPP) substrate system (N2765; Sigma-Aldrich) as previously described[38]. MtVs and standard curve were incubated with the pNPP substrate at 37°C for 5 min. Absorbance was measured at 405 nm with 655 nm as a reference wavelength.

#### 2.6.5. MtV-Collagen Calcification Assay

To assess the calcification capacity of the MtVs, we assessed their mineralization in cell-free conditions following previously reported protocols[39]. Briefly, tissue culture plates were coated with type I collagen (Corning) in a 0.01%(w/v) solution in 0.1 M acetic acid at room temperature for 4 h. Following coating, 50 μg/ml of MtVs were applied to the collagen coated surfaces and left overnight in 4°C. Samples were then exposed to mineralization medium (α-MEM with 15% FBS and 10 mM β-glycerophosphate) to create an acellular MtV-collagen culture which was incubated at 37°C. Medium was changed three times a week for a total of 2 weeks. The MtV-functionalized collagen model was used to assess the MtV calcium phosphate content without exposure to mineralization medium. MtV-free collagen surfaces (PBS alone) were used as the untreated control.

### 2.7. Cell Uptake Assay for EVs

The uptake of MtVs was evaluated as previously described[40]. Briefly, vesicles were labelled with Cell Mask™ Deep Red Plasma Membrane Stain (5 μg/ml in PBS) (Thermo Scientific) for 10 min, then washed twice with PBS via ultracentrifugation at 120,000 g for 70 min. hBMSCs were seeded at a density of 4 × 10^3^ cells/cm^2^. After 24 h, the medium was replaced with fresh basal medium containing labelled MtVs. After 8 h incubation, cells were fixed with neutral buffered formalin, permeabilized with triton-X, the actin skeleton labelled with Alexa Fluor 488 phalloidin (1:20) (Cell Signalling Technology) and then mounted and coverslipped with Prolong ™ Gold Antifade Mountant with DAPI (Thermo Scientific) to label the nuclei. Samples were imaged with the aforementioned EVOS fluorescence inverted microscope.

### 2.8. hBMSCs Proliferation and Migration

The effects of MtVs on hBMSC proliferation was assessed by quantifying DNA content. Cells were seeded at 3 × 10^3^ cells/cm^2^ in basal media for 24 h. Media was replaced with fresh basal medium supplemented with MtVs (5 µg/ml). Cells cultured in basal medium alone were used as a control. DNA content was evaluated on day 3 and 7 utilising the previously described DNA quantification assay.

Migration rate was assessed by conducting the scratch assay as previously described[41]. Briefly, cells at 30 x 10^3^ cells/cm^2^ were plated in a 48-well plate in basal medium. A scratch was generated with a 200 µl tip and the baseline width was measured. Cells were then incubated with basal medium containing 5 µg/ml MtVs for 3 days. Cells culture in basal medium alone was used as the control. Migration rate calculated as percentage wound area closure from between day 0 and day 3 was evaluated under a Leica DMi1 microscope (Leica).

### 2.9. MtV-induced hBMSC Osteogenesis

Cells were seeded in 96-well plates at a density of 21 × 10^3^ cells/cm^2^ in basal medium. After 24 h, the medium was replaced with mineralization medium (consisting of basal medium with 10 mM β-glycerophosphate and 10 nM dexamethasone) supplemented with MtVs (5 µg/ml) for 14 days, medium was changed every 2 days. Cells cultured in mineralization medium alone were used as the control.

### 2.10. hBMSCs ALP Activity

ALP activity of MtVs-treated hBMSCs was evaluated using the p-nitrophenyl phosphate (pNPP) substrate system (N2765; Sigma-Aldrich) as previously described[38]. Cells were lysed with Triton X-100 (0.1%, Sigma-Aldrich) with several freeze/thaw cycles. Cell lysate and standard curve were incubated with the pNPP substrate at 37°C for 5 min. Absorbance was measured at 405 nm with 655 nm as a reference wavelength. ALP activity was normalized with DNA content using the Quant-iT Picogreen dsDNA assay (P11496, Thermo Fisher Scientific). The ALP activity of MtVs added during culture was quantified and used for normalization.

### 2.11. Calcium Deposition and Mineralization

Alizarin red S staining was conducted to visualize calcium deposits as previously described[41]. Briefly, samples were washed with PBS and fixed in 10% NBF for 30 min. Following which, samples were washed with distilled water and incubated with 40 mM Alizarin Red S solution (pH 4.1, Sigma Aldrich) for 10 min. The unbound staining solution was removed by distilled water washes. Staining was visualized using light microscopy (Olympus Microscope BX43). For alizarin red quantification, samples were de-stained with 10% cetylpyridinium chloride (Sigma-Aldrich) for 1 h and the absorbance was read at 550 nm using the CLARIOstar Plus microplate reader (BMG Labtech).

To detect mineral deposits, von Kossa staining was performed by incubating the sections with 1% silver nitrate (209 139, Sigma-Aldrich) directly under a light bulb (Philips Master TL5HO 54W 830), for 1 h. The samples were subsequently washed with 5% sodium thiosulfate (A17629, Alta Aesar) and imaged using light microscopy (Olympus Microscope BX43). The mean mineral nodule percentage coverage was quantified using ImageJ (NIH).

### 2.12. Angiogenic Analysis

Human endothelial colony forming cells (hECFCs) were expanded to 80% confluence on tissue culture polystyrene, coated with 0.1% gelatin. For the tube formation assay, μ-slides for angiogenesis (Ibidi, Gräfelfing) were coated with growth factor-reduced Matrigel (1:1 diluted with PBS, 45 min at 37°C, Corning). hECFCs (48,000 cell/cm^2^) were seeded in EBM-2 supplemented with 1% FBS and MtVs (10 µg/ml), using 6 replicates per condition. EV-free medium was used as the control. Slides were incubated for 8 h at 37°C under humidified conditions and with 5% CO_2_, followed by observation of tube networks and image capture by microscopy (Olympus Microscope BX43). Tube network quantification was conducted using the “Angiogenesis Analyzer” plugin for ImageJ (NIH).

### 2.13. BMP2 and VEGF Quantification

MtVs were lysed with Triton X-100 (1%, Sigma-Aldrich) with several freeze/thaw cycles. The BMP2 and VEGF content of MtVs was quantified using the R&D Systems DuoSet® enzyme-linked immunosorbent assay (ELISA) according to the manufacturer’s protocol.

### 2.14. LC-MS Sample Preparation and Proteomics Analysis

Three independent sample preparations of HB MtVs were assessed using a label-free MS-LC/LC approach. EV proteins were denatured and alkylated in 50 µl 8 M Urea, 1 M ammonium bicarbonate (ABC) containing 10 mM TCEP (tris (2-carboxyethyl) phosphine hydrochloride) and 40 mM 2-chloro-acetamide for 30 min. After 8-fold dilution with 100mM HEPES (pH8), 10 mM CaCl, Proteins were digested overnight with Trypsin/LysC mixture (1 µg) (Thermo). Peptides were desalted with homemade C-18 stage tips (3 M, St Paul, MN), eluted with 80% Acetonitrile (ACN) and, after evaporation of the solvent in the speedvac, redissolved in buffer A (0,1% formic acid). Peptides were separated on a 30 cm pico-tip column (75 µm ID, New Objective) in-house packed with 1.9 µm aquapur gold C-18 material (dr. Maisch) using a 120 min gradient (7% to 80% ACN 0.1% FA), delivered by an Vanquish Neo HPLC (Thermo), and electro-sprayed directly into a Orbitrap Eclipse Tribrid Mass Spectrometer (Thermo Scientific). The latter was set in data dependent mode with a cycle time of 1.2 second for both Faims CV settings (−45V and −65V), in which the full scan over the 400-1200 mass range was performed at a resolution of 240K. The most intense ions (charge state 2-6) were isolated by the quadrupole with a 0.4 Da window and fragmented with a HCD collision energy of 30% and analysed in the iontrap. The maximum injection time of the ion trap was set to 35 milliseconds. Dynamic exclusion of 10 ppm was set on 45 seconds, excluding isotopes. Raw files were analyzed with the Maxquant software (version 2.6.7.0) with oxidation of methionine set as variable modifications, and carbamidomethylation of cysteine set as fixed modification. The Human protein database of Uniprot (2024) was searched with both the peptide as well as the protein false discovery rate set to 1%. To determine proteins of interest, the protein groups output file was used to perform a differential enrichment analysis.

The use of stringent criteria only permitted the inclusion of proteins identified in a least two biological replicates, with > 2 spectral counts in at least one repeat. The protein annotation through evolutionary relationship (PANTHER) classification systems (version 19.0) was used for gene ontology (GO) annotation of biological pathways, molecular mechanisms and cellular components of proteins found in HB MtVs and their associated Kyoto Encyclopaedia of Genes and Genomes (KEGG) pathways. GO and KEGG pathway results were filtered for those involved in bone tissue. StringDB was used to generate a protein-protein interaction network.

### 2.15. Statistics

For all data presented, experiments were repeated at least 3 times. All statistical analysis was undertaken using ANOVA multiple comparisons test with Tukey modification (GraphPad Prism 8). P values equal to or lower than 0.05 were considered significant. *P ≤ 0.05, **P ≤ 0.01 and ***P ≤ 0.001.

## 3. Results

### 3.1. Chondrogenic differentiation of hBMSC microtissues

Histological analysis was conducted to evaluate the effects of differing culture conditions on hBMSC chondrogenesis (Figure 2). Assessment of chondrogenic markers (Safranin O, Toluidine Blue, collagen type II) showed the BMP2 addition in both the chondrogenic and hypertrophic medium conditions increased the intensity and uniformity of GAG presence and collagen type II deposition throughout the microtissue when compared to the BMP2-free groups, which displayed weak GAG and collagen type II staining at the periphery of the microtissues. The BMP2 groups exhibited increased staining intensity for markers of hypertrophy (collagen type I, X, VEGF, ALP) when compared to the respective BMP2-free microtissues, with hypertrophic groups displaying enhanced expression of especially ALP and collagen type I compared to the chondrogenic conditions. Moreover, microtissues which exhibited a more hypertrophic phenotype tend to display enlarged lacunae when compared to chondrogenic groups. Additionally, the extent of microtissue matrix mineralization was assessed via Alizarin red and von Kossa staining for calcium deposits and mineralization, respectively (Figure S1). Our findings showed a similar degree of calcium accumulation across all microtissue groups, with weak staining primarily situated within the cells. Moreover, all groups displayed negligible von Kossa staining for mineral deposits.

**Figure 2.**
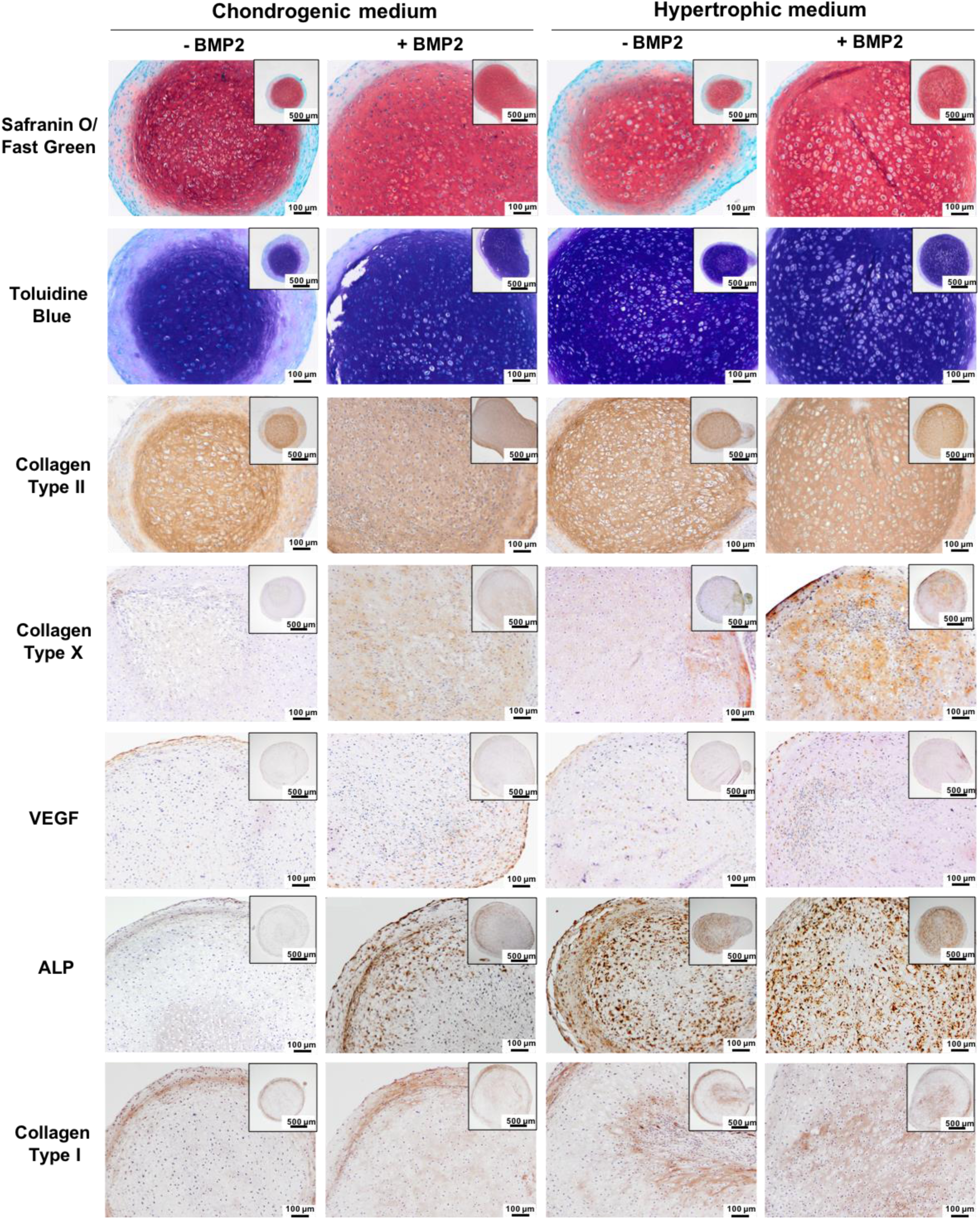
*In vitro* chondrogenesis of hBMSC microtissues differentiated with or without BMP2 in chondrogenic or hypertrophic conditions. Sections of cartilaginous microtissues from different culture conditions (+/- BMP2) (chondrogenic/hypertrophic medium) were stained for GAGs (Safranin O and Toluidine blue staining), collagen type I, collagen type II, collagen type X, VEGF and ALP after day 28 of culture.

Additionally, the influence of culture condition on microtissue size was assessed. In both chondrogenic and hypertrophic groups, the addition of BMP2 significantly increased microtissue diameter when compared to BMP2-free microtissues (1.13-, 1.11-fold, respectively) (P ≤ 0.01) (Figure 3A). These findings correlated with similar upregulation in microtissue total protein content induced by BMP2 conditioning (Figure 3B), despite similar DNA content across groups (Figure 3C). Moreover, the BMP2 chondrogenic culture group exhibited significantly increased GAG content when compared to the BMP2-free chondrogenic group (P ≤ 0.01) (Figure 3D).

**Figure 3.**
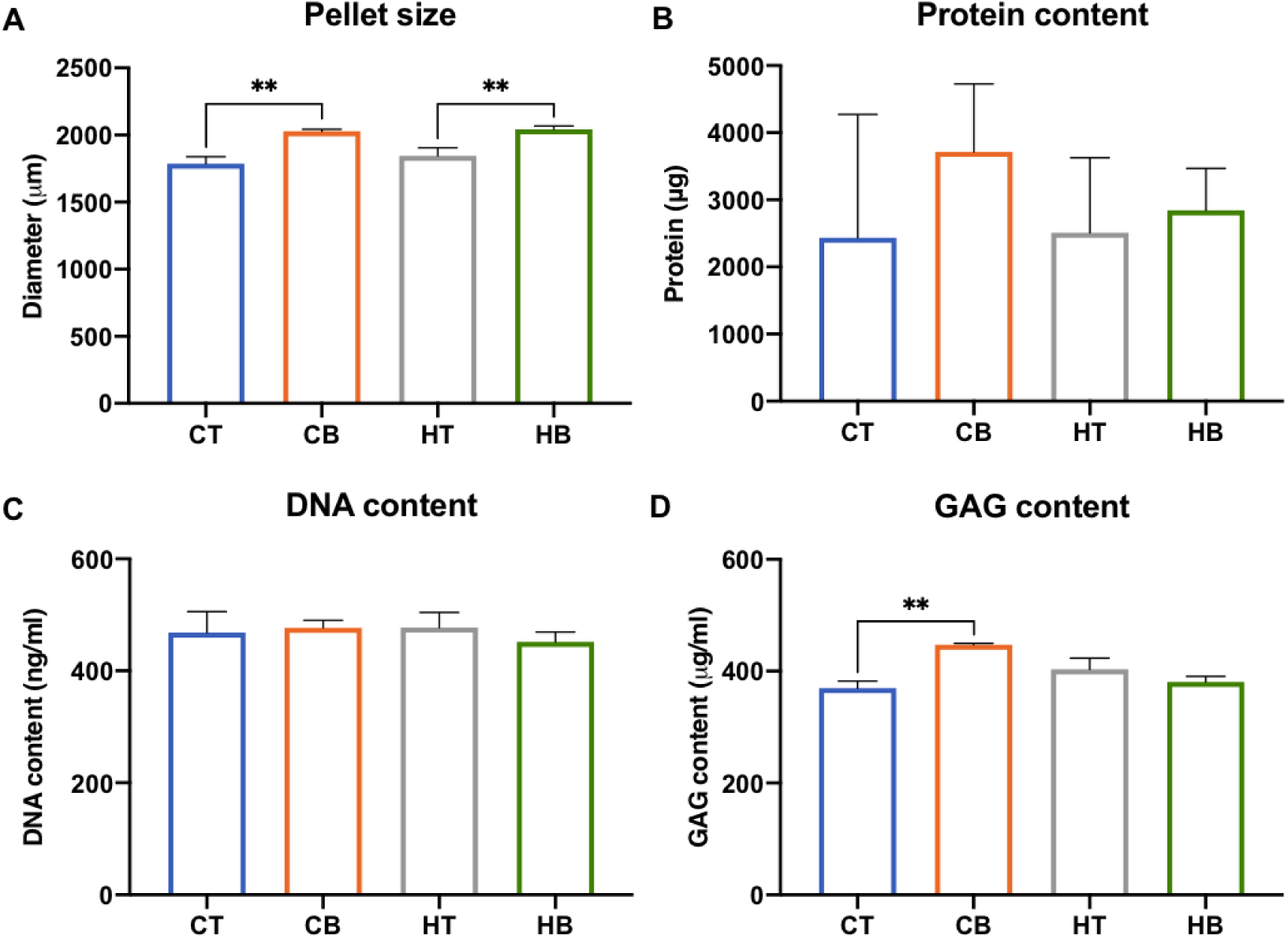
Biochemical analysis of cartilaginous microtissue A) size, B) protein, C) DNA and D) GAG content after 28 days of culture. Data expressed as mean ± SD (N = 3). **P ≤ 0.01. CT = chondrogenic medium/-BMP2; CB = chondrogenic medium/+BMP2; HT = hypertrophic medium/-BMP2; HB = hypertrophic medium/+BMP2.

### 3.2. Characterization of MtVs

Following 28 days of culture, MtVs were obtained through collagenase type II digestion and differential ultracentrifugation. Leveraging size-dependent sedimentation via differential ultracentrifugation, two MtVs subpopulations were obtained from the microtissues based on size. TEM imaging showed the L-MtVs and S-MtVs subpopulations exhibited a spherical morphology, typical for these nano-sized particles (Figure 4A). NTA analysis demonstrated that the vesicles displayed an average diameter of 157 and 74 nm for the L-MtVs and S-MtVs, respectively (Figure 4B). Moreover, there was a substantially higher concentration of S-MtVs particles (2.92 ± 0.09 x 10^9^/ml) when compared to the L-MtVs (1.31 ± 0.25 x 10^9^/ml). Quantification of MtV protein content showed that the addition of BMP2 significantly improved MtV yield when compared to nanoparticles derived from the BMP2-free microtissues, with hypertrophic conditioning further improving nanoparticle yield for L-MtVs (Figure 4C). Both MtV populations exhibited positive CD63 tetraspanin marker expression, with the S-MtVs displaying a 1.18 - 1.89-fold increase in CD63 content compared to the L-MtVs (Figure 4D). Within the S-MtV population, there was substantial increase in CD63 positive particles from the BMP2 treated microtissues. Further, the influence of BMP2/hypertrophic conditioning on the microtissue secretome was evaluated at days 21 and 28. The quantity of proteins released into the medium was significantly reduced in the BMP2-cultured microtissues at day 21 and day 28 (Figure S2A), while hypertrophic conditioning enhanced the secretrome protein content when compared to the chondrogenic groups at day 28. In contrast, the proportion of EVs within the secretome were significantly elevated by BMP2 treatment and hypertrophic induction at day 28 (Figure S2B, S2C). As MtVs are known to be initiators of mineral deposition, we investigated the influence of different culture conditions on the ALP activity of these nanoparticles (Figure 4E). Our findings showed that all MtV groups exhibited positive ALP activity, with BMP2 conditioning substantially improving activity when compared to the MtVs derived from the respective BMP2-free groups. Moreover, exposure to hypertrophic medium further elevated MtVs’ ALP activity in both BMP2 and BMP2-free groups when compared to microtissues cultured in chondrogenic medium.

**Figure 4.**
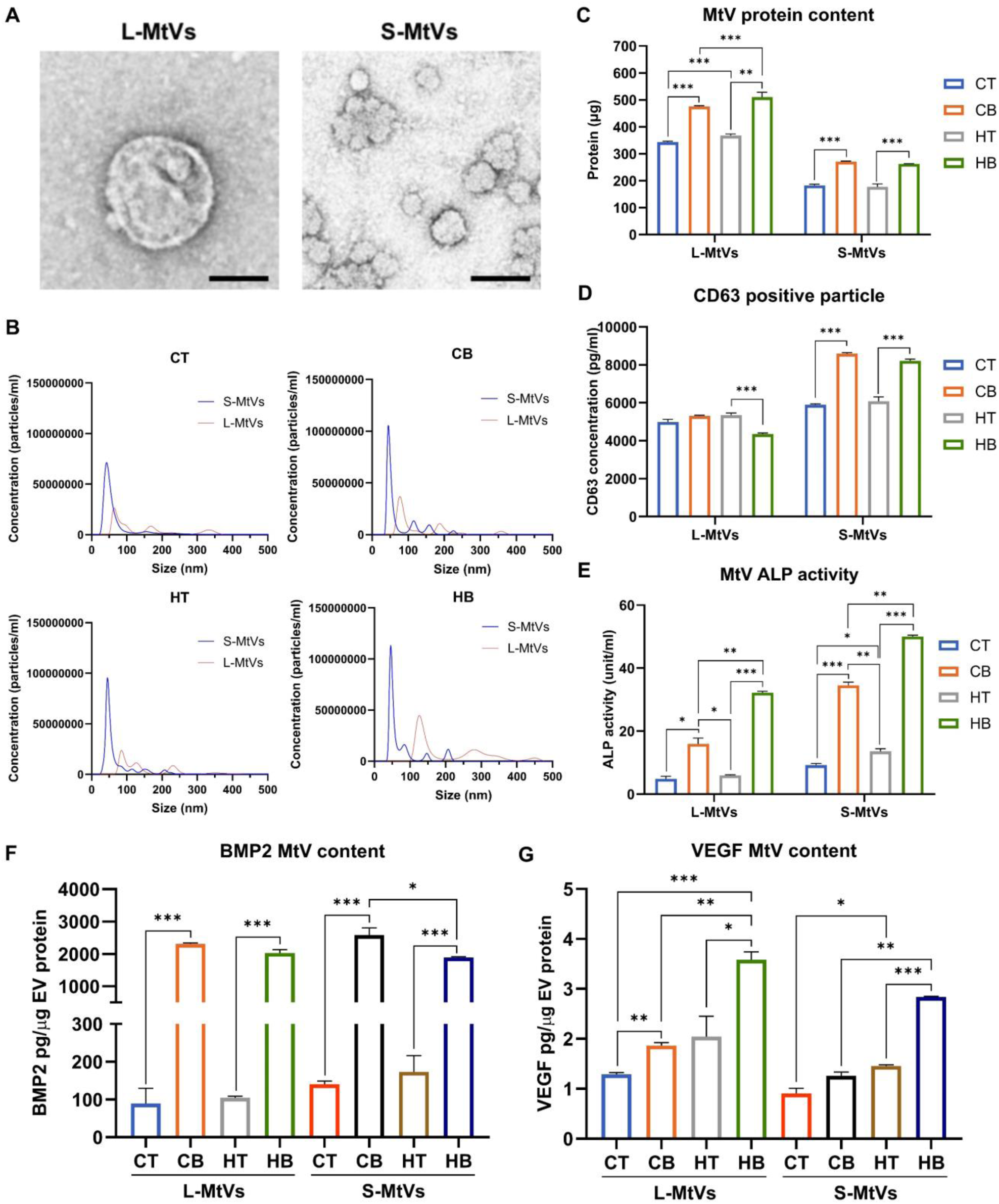
Characterization of cartilaginous microtissue-derived MtVs. A) Representative TEM image of L-MtVs and S-MtVs. Scale bar = 100 nm. B) NTA of isolated MtVs. C) MtV protein quantification. D) Quantification of CD63-positive particles E) ALP activity of MtVs. Quantification of A) BMP2 and B) VEGF content within MtVs measured via R&D Systems DuoSet® ELISA. BMP2 and VEGF contents were normalized to EV protein content. Data expressed as mean ± SD (N = 3). *P ≤ 0.05, **P ≤ 0.01 and ***P ≤ 0.001. CT = chondrogenic medium/-BMP2; CB = chondrogenic medium/+BMP2; HT = hypertrophic medium/-BMP2; HB = hypertrophic medium/+BMP2.

### 3.3. Quantification of MtV-associated bioactive factors

To determine whether the soft callus-mimetic MtVs were enriched with pro-osteogenic/angiogenic growth factors, by quantifying BMP2 and VEGF content. Our findings showed that hypertrophic conditioning slightly increased MtV BMP2 levels when compared to vesicles from chondrogenic conditions (Figure 4F) (P > 0.05). BMP2 treatment significantly enhanced the quantity of BMP2 found within the MtVs compared to the BMP2-free groups (P ≤ 0.001). In S-MtVs, hypertrophic conditioning in conjunction with BMP2 treatment reduced BMP2 levels when compared to BMP2 and chondrogenic culture. Similarly, the VEGF content of these MtVs were slightly elevated in the groups treated with either BMP2 or hypertrophic medium when compared to BMP2-free and chondrogenic conditions respectively (Figure 4G) (P ≤ 0.05 - 0.001). Moreover, synergistic BMP2 and hypertrophic conditioning further enhanced VEGF content within both L-MtVs and S-MtVs when compared to groups conditioned with only BMP2 or hypertrophy (P ≤ 0.05 - 0.01).

### 3.4. MtVs stimulate the calcification of collagen in acellular conditions

As MtVs are known to be initiators of ECM mineral nucleation, we evaluated the biomineralization capacity of these nanoparticles in acellular conditions (Figure 5). Following the binding of MtVs to collagen-coated surfaces, samples were kept in mineralization medium for 2 weeks. Our findings showed that MtVs from chondrogenic groups with no BMP2 did not improve calcium accumulation within the collagen-coated surfaces, showing similar amounts as the EV-free control. In contrast, MtVs derived from BMP2 and/or hypertrophic conditioned microtissues substantially increased calcium deposition when compared to EV-free control (>5-fold) and the microtissues cultured in BMP2-free (>1.6-fold) or chondrogenic conditions (2.55-fold). Moreover, the S-MtV subpopulation exhibited substantially enhanced calcium accumulation (2.31 - 6.9-fold) when compared to their respective L-MtV group. Alizarin red staining was conducted on MtV-functionalized collagen surfaces to determine whether the MtVs contained calcium phosphate (Figure S3). Our finding showed that all MtV groups exhibited negligible staining for calcium content, consistent with the MtV-free control (P> 0.05).

**Figure 5.**
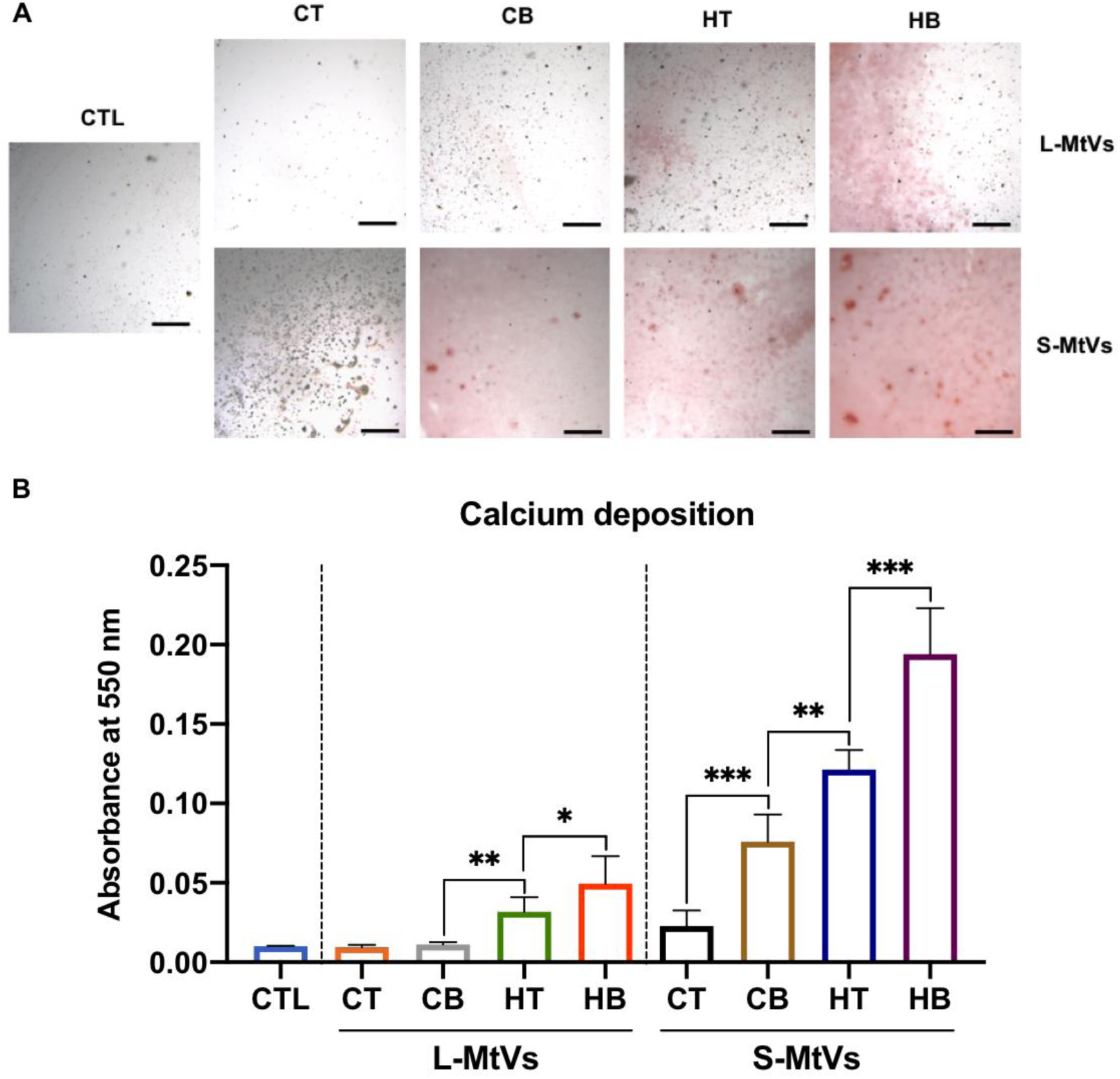
The mineralization capacity of MtVs in acellular conditions. A) Alizarin red S staining for calcium deposition of MtV-functionalized collagen surfaces. Scale bar = 200 µm. B) Semi-quantification of calcium deposition. Data expressed as mean ± SD (N = 3). *P ≤ 0.05, **P ≤ 0.01 and ***P ≤ 0.001. CTL = control; CT = chondrogenic medium/-BMP2; CB = chondrogenic medium/+BMP2; HT = hypertrophic medium/-BMP2; HB = hypertrophic medium/+BMP2.

### 3.5. The effect of MtVs on hBMSC uptake, proliferation and migration

Following the characterization of these MtVs, we evaluated their biological function by initially assessing their capacity to be taken up by recipient hBMSCs (Figure 6A). After 8 hours of incubation, we observed that the L-MtVs and S-MtVs were internalized by the recipient hBMSCs for all groups. Untreated control samples are shown in supplementary Figure S4. Moreover, we evaluated the influence of MtV treatment on stimulating hBMSC proliferation (Figure 6B). Our results showed that the MtV treatment exhibited a time-dependent increase in hBMSCs proliferation when compared to the untreated control. Moreover, it was observed that MtVs derived from BMP2-treated microtissues improved the proliferative capacity of treated hBMSCs when compared to the MtVs procured from the BMP2-free groups. Additionally, treatment with MtVs of each groups improved the migration of hBMSCs when compared to the untreated group (Figure 6C). Notably, in both L-MtVs and S-MtVs subpopulations, vesicles derived from microtissues exposed to BMP2 stimulation showed an increased ability to accelerate hBMSC migration rate compared to those from BMP2-free microtissues. Similarly, vesicles from hypertrophic microtissues demonstrated enhanced migration-promoting effects relative to those from chondrogenic microtissues. Importantly, when both BMP2 stimulation and hypertrophic conditions were combined, the vesicles exhibited an even more pronounced effect on accelerating hBMSC migration, suggesting a synergistic enhancement of their migration-promoting properties.

**Figure 6.**
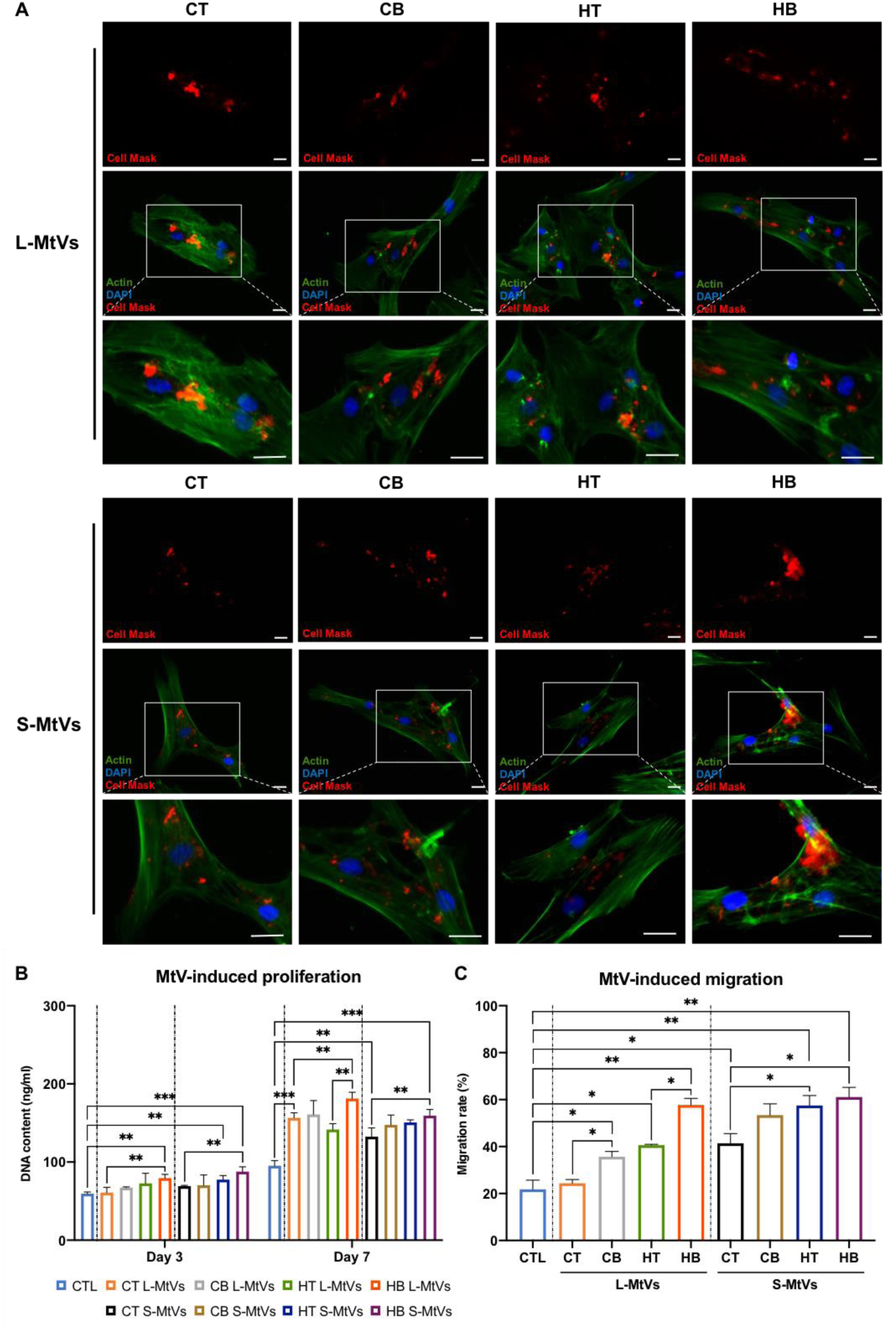
The general effects of MtVs on recipient hBMSCs. A) DAPI/f-actin/Cell Mask staining showing MtV uptake by hBMSCs following 8 hours incubation. Scale bar = 10 µm. B, C) The effects of MtV treatment on hBMSC (B) proliferation and (C) migration. Data expressed as mean ± SD (N = 3). *P ≤ 0.05, **P ≤ 0.01 and ***P ≤ 0.001. CTL = control; CT = chondrogenic medium/-BMP2; CB = chondrogenic medium/+BMP2; HT = hypertrophic medium/-BMP2; HB = hypertrophic medium/+BMP2.

### 3.6. MtVs enhance the mineralization capacity of hBMSCs

The effects of MtV treatment on hBMSC osteogenic differentiation and mineralization was assessed by quantifying ALP activity, calcium production and mineral nodule formation following 14 days in osteogenic culture (Figure 7, Figure S5). ALP activity was substantially increased in hBMSCs treated with all MtV groups (1.15 - 3.12-fold) when compared to the untreated control after 7 days of osteogenic culture (Figure S5). In particular, we observed enhanced ALP activity in hBMSCs treated with MtVs obtained from BMP2 and hypertrophic conditioned microtissues compared to vesicles from BMP2-free and chondrogenic groups. Moreover, our findings showed that all MtV treated groups exhibited significantly increased hBMSC mineralization when compared to the EV-free control confirmed through alizarin red staining for calcium deposition (2.78 - 12.97-fold) and von Kossa staining for mineral nodules (6.02 - 12.77-fold) (Figure 7A). Interestingly, MtVs derived from BMP2-conditioned microtissues, cultured in either chondrogenic or hypertrophic medium, substantially improved hBMSC osteogenic differentiation when compared to MtVs from the BMP2-free groups (Figure 7B). Similarly, MtVs from the hypertrophic microtissues significantly promoted mineralization when compared to the chondrogenic groups, whilst the S-MtV subpopulation exhibited enhanced osteoinductivity when compared to the L-MtVs.

**Figure 7.**
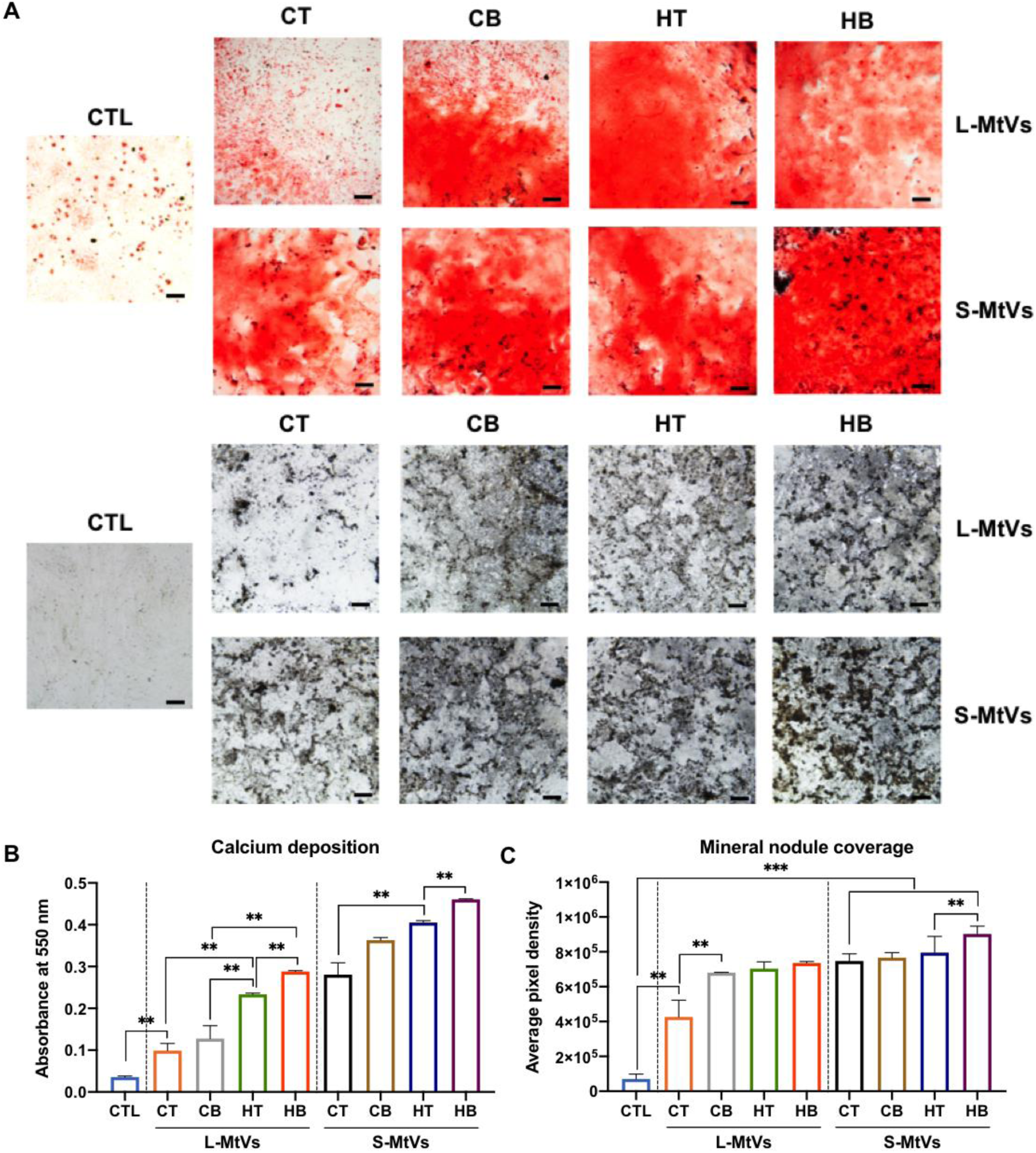
The effects of MtVs on hBMSC osteogenic differentiation. A) Alizarin red S staining for calcium deposition, B) Von Kossa staining for mineral nodules. Scale bars = 200 µm. Semi-quantification of C) calcium deposition and D) mineral nodule coverage. Data expressed as mean ± SD (N = 3). *P ≤ 0.05, **P ≤ 0.01 and ***P ≤ 0.001. CTL = control; CT = chondrogenic medium/-BMP2; CB = chondrogenic medium/+BMP2; HT = hypertrophic medium/-BMP2; HB = hypertrophic medium/+BMP2.

### 3.7. Soft callus-mimetic MtVs improve the angiogenic tube formation of hECFCs

In addition to assessing the osteogenic potency of soft callus-mimetic MtVs, we evaluated the capacity of the MtVs to promote hECFC angiogenic tube formation (Figure 8). Our findings showed that MtV treatment improved the tube formation potential of hECFCs when compared to the untreated control (i.e. number of nodes, junctions, and total tube length), with MtVs from the BMP2 and hypertrophic microtissues stimulating further enhanced hECFC angiogenesis.

**Figure 8.**
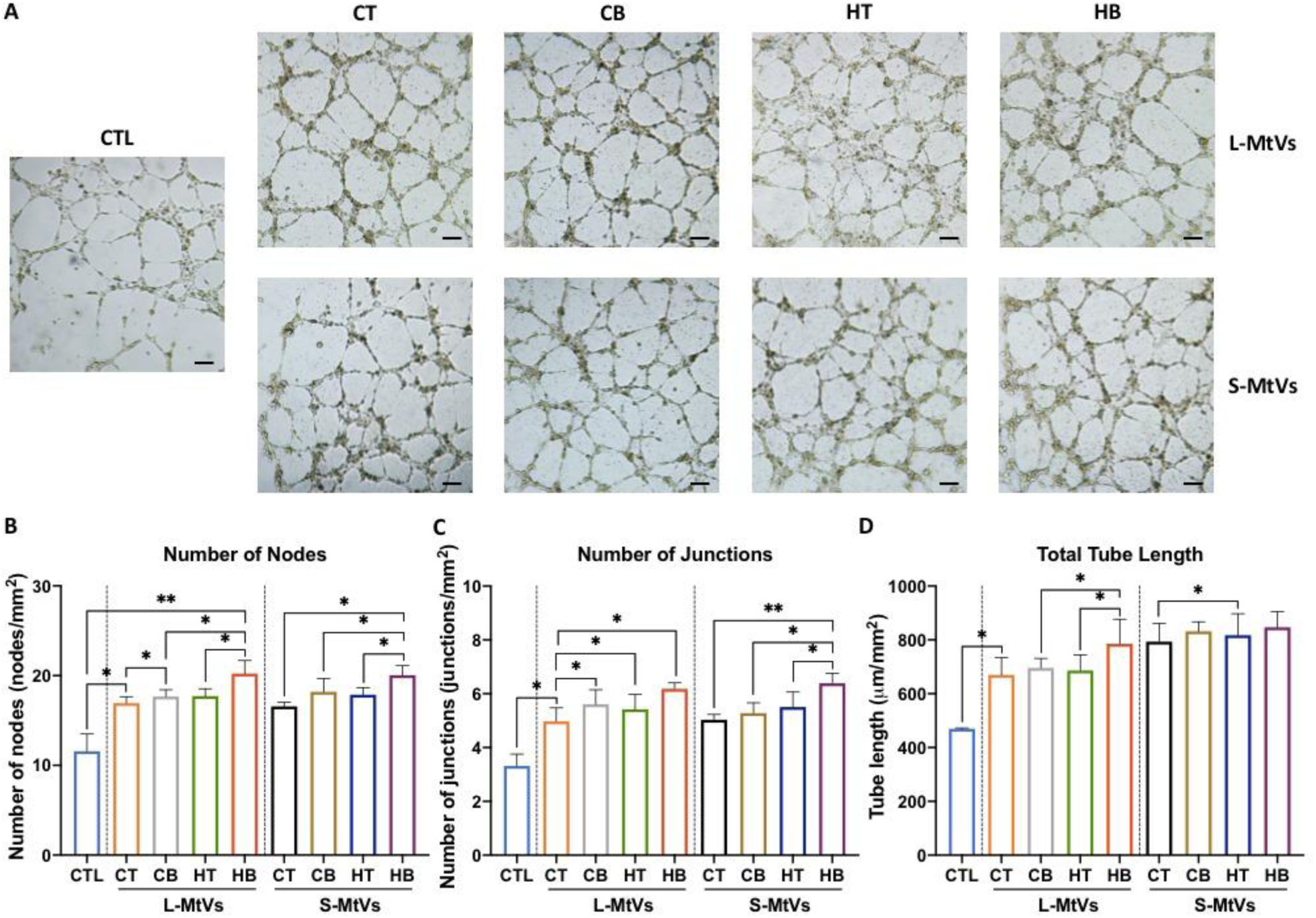
MtV-induced angiogenesis of hECFCs. A) Representative microscopy images of endothelial networks formed by hECFCs after 8 h incubation with MtVs. Scale bar = 200 µm. Results of hECFC network quantification measuring, B) number of nodes, C) number of junctions, and D) tube length per mm^2^. Data expressed as mean ± SD (N = 3). *P ≤ 0.05, **P ≤ 0.01 and ***P ≤ 0.001. CTL = control; CT = chondrogenic medium/-BMP2; CB = chondrogenic medium/+BMP2; HT = hypertrophic medium/-BMP2; HB = hypertrophic medium/+BMP2.

### 3.8. The proteome of HB-MtVs is enriched in proteins involved in ECM organization and ossification

Having demonstrated the HB MtVs exhibited the most potent osteoinductive and angiogenic capabilities, the functional landscape of MtVs associated proteins were characterized by performing proteomic analysis followed by GO and KEGG pathway enrichment analysis. GO analysis revealed that HB MtV-associated proteins were enriched for biological processes linked to bone ECM mineralization, osteogenesis and regeneration, specifically through the enrichment of organophosphate metabolic process, and phosphorus metabolic process, bone trabecula formation, regulation of Wnt signalling pathway, all of which are fundamental for bone formation (Figure 9A). In the cellular component category, many of the proteins localized to the EVs and the extracellular space, confirming the vesicular and secretory origin of the MtVs and their involvement in ECM organization (Figure 9B). Molecular function analysis indicated enrichment in calcium ion binding, ECM structural constituents, and ECM remodelling enzymes, all of which are critical for cartilage matrix remodelling and mineral deposition (Figure 9C). KEGG pathway enrichment analysis further emphasized the involvement of HB-MtV proteins in biological signalling pathways central to skeletal development, including Integrin signalling (87 genes), VEGF signalling (28 genes), Angiogenesis (62 genes), PI3K-Akt signalling (24 genes), and Wnt signalling (44 genes) (Figure 9D). These pathways are closely associated with EO and chondrocyte hypertrophy, reflecting the functional phenotype of the originating tissue. Additional pathways, including FGF, Ras, cytoskeletal regulation by Rho GTPase, cadherin, TGF-beta, underscore the mechanistic diversity by which MtVs influence matrix integrity and structure. Collectively, these findings indicate that MtVs derived from hypertrophic cartilage microtissues are enriched in proteins involved in matrix remodelling, cell-matrix communication, and mineralization, supporting their proposed role in skeletal development.

**Figure 9.**
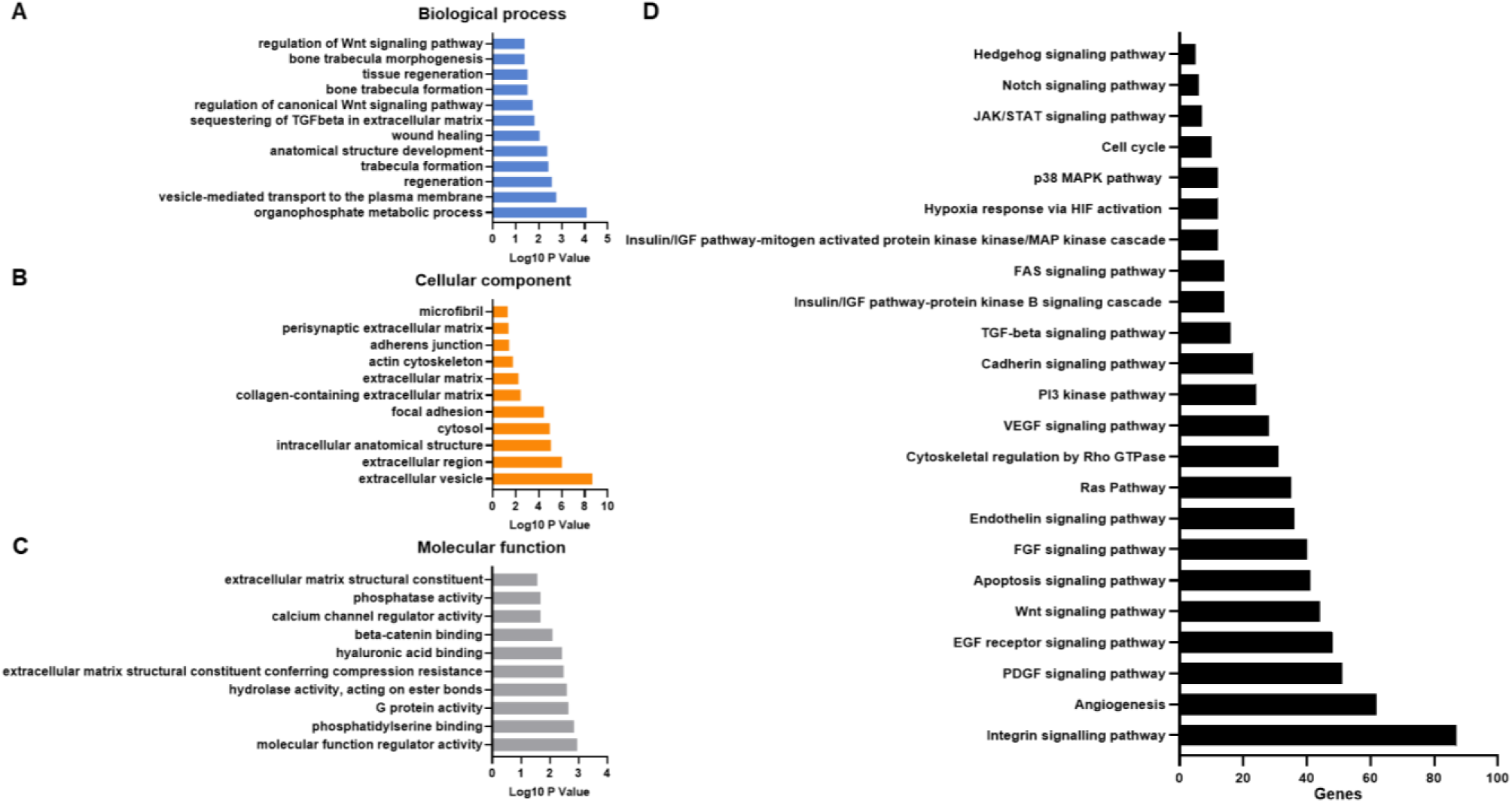
Gene ontology analysis of proteins within HB-MtVs. Top ten GO prediction scores covering the domains of A) biological processes, B) cellular compartments and C) molecular mechanisms of HB-MtVs associated proteins. D) KEGG pathway analysis.

Table 1 details individual HB MtV proteins directly involved in bone formation and matrix regulation. Key proteins such as fibronectin (FN1), collagen type I (COL1A1, COL1A2), osteopontin (SPP1), alkaline phosphatase (ALPL), and annexins (ANXA2, ANXA5) were identified, each linking to crucial matrix and mineralization pathways. Regulatory proteins, including RUNX2, BMP2, β-catenin (CTNNB1), and MAPK14, further illustrate a coordinated molecular signature supporting osteogenesis. To better understand how these bone-related proteins may interact, a protein network was constructed using String DB and is shown in Figure 10. This network analysis identified distinct clusters with high connectivity, including a dense core of ribosomal/translational machinery proteins (RPL, RPS, EIF) suggesting coordinated roles in translation and potentially ECM remodelling. Another prominent cluster enriched for ECM proteins (collagens, FN1, SPARC, BGN) supports matrix organization, while COL10A1 highlights the hypertrophic chondrocyte origin. The network also featured angiogenesis-associated proteins (TGFβ1, ENG, SPARC, THBS1), vesicle trafficking and immune modulators (ANXA5, VIM, S100A10), and metabolic proteins (ATP5F1B, PKM, LDHA), indicating that HB MtVs may synergistically support cellular translation, matrix maturation, vascular invasion, immune crosstalk, and local energy metabolism.

**Figure 10.**
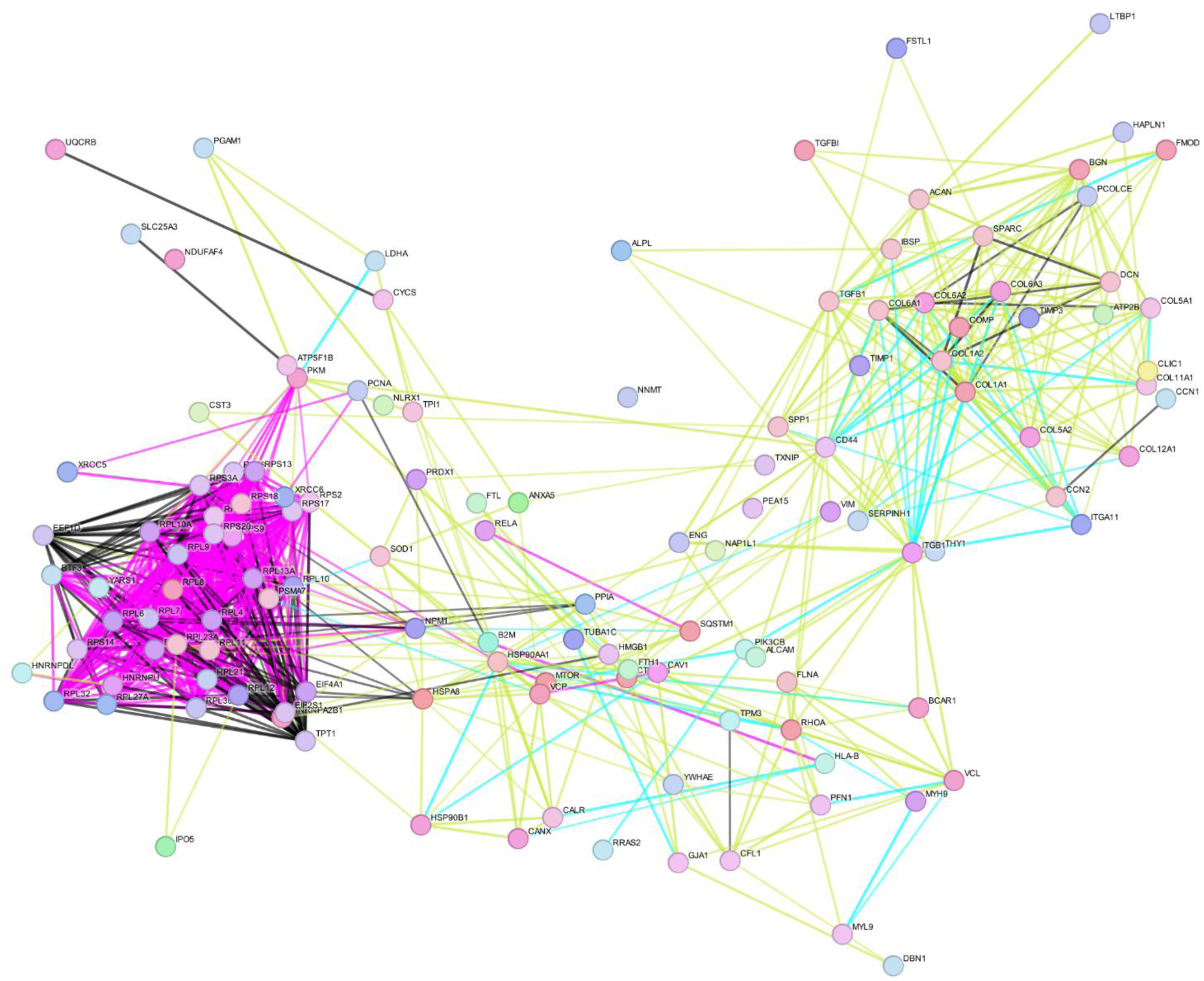
STRING network analysis of bone-related proteins identified within HB MtVs. Nodes represent individual proteins and are color-coded by functional cluster. Lines between nodes indicate protein associations, color-coded by evidence type: black for co-expression, cyan for curated database, yellow for text-mining, and magenta for experimental evidence.

**Table 1.**
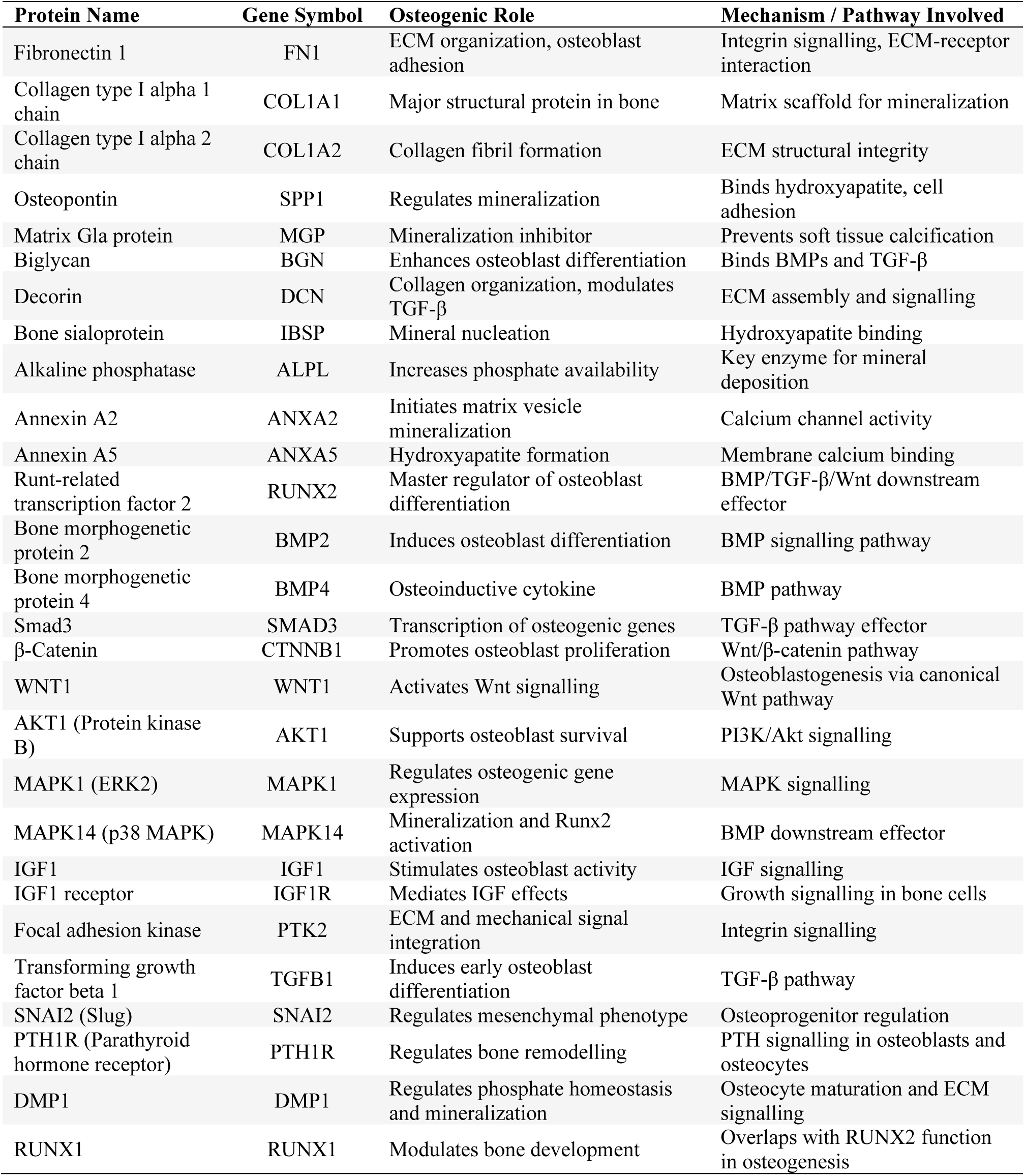
HB MtV proteins related to osteogenic mechanisms.

| Protein Name | Gene Symbol | Osteogenic Role | Mechanism / Pathway Involved |
| --- | --- | --- | --- |
| Fibronectin 1 | FN1 | ECM organization, osteoblast adhesion | Integrin signalling, ECM-receptor interaction |
| Collagen type I alpha 1 chain | COL1A1 | Major structural protein in bone | Matrix scaffold for mineralization |
| Collagen type I alpha 2 chain | COL1A2 | Collagen fibril formation | ECM structural integrity |
| Osteopontin | SPP1 | Regulates mineralization | Binds hydroxyapatite, cell adhesion |
| Matrix Gla protein | MGP | Mineralization inhibitor | Prevents soft tissue calcification |
| Biglycan | BGN | Enhances osteoblast differentiation | Binds BMPs and TGF-β |
| Decorin | DCN | Collagen organization, modulates TGF-β | ECM assembly and signalling |
| Bone sialoprotein | IBSP | Mineral nucleation | Hydroxyapatite binding |
| Alkaline phosphatase | ALPL | Increases phosphate availability | Key enzyme for mineral deposition |
| Annexin A2 | ANXA2 | Initiates matrix vesicle mineralization | Calcium channel activity |
| Annexin A5 | ANXA5 | Hydroxyapatite formation | Membrane calcium binding |
| Runt-related transcription factor 2 | RUNX2 | Master regulator of osteoblast differentiation | BMP/TGF-β/Wnt downstream effector |
| Bone morphogenetic protein 2 | BMP2 | Induces osteoblast differentiation | BMP signalling pathway |
| Bone morphogenetic protein 4 | BMP4 | Osteoinductive cytokine | BMP pathway |
| Smad3 | SMAD3 | Transcription of osteogenic genes | TGF-β pathway effector |
| β-Catenin | CTNNB1 | Promotes osteoblast proliferation | Wnt/β-catenin pathway |
| WNT1 | WNT1 | Activates Wnt signalling | Osteoblastogenesis via canonical Wnt pathway |
| AKT1 (Protein kinase B) | AKT1 | Supports osteoblast survival | PI3K/Akt signalling |
| MAPK1 (ERK2) | MAPK1 | Regulates osteogenic gene expression | MAPK signalling |
| MAPK14 (p38 MAPK) | MAPK14 | Mineralization and Runx2 activation | BMP downstream effector |
| IGF1 | IGF1 | Stimulates osteoblast activity | IGF signalling |
| IGF1 receptor | IGF1R | Mediates IGF effects | Growth signalling in bone cells |
| Focal adhesion kinase | PTK2 | ECM and mechanical signal integration | Integrin signalling |
| Transforming growth factor beta 1 | TGFB1 | Induces early osteoblast differentiation | TGF-β pathway |
| SNAI2 (Slug) | SNAI2 | Regulates mesenchymal phenotype | Osteoprogenitor regulation |
| PTH1R (Parathyroid hormone receptor) | PTH1R | Regulates bone remodelling | PTH signalling in osteoblasts and osteocytes |
| DMP1 | DMP1 | Regulates phosphate homeostasis and mineralization | Osteocyte maturation and ECM signalling |
| RUNX1 | RUNX1 | Modulates bone development | Overlaps with RUNX2 function in osteogenesis |

## 4. Discussion

Due to the intrinsic role of EVs in tissue development and homeostasis, there has been tremendous interest in elucidating the function of these nanoparticles in several biological processes, with the potential to harness these vesicles for regenerative medicine[42, 43]. EVs have been shown to carry multiple bioactive growth factors, important for orchestrating the spatially and temporally complex processes involved in EO[31, 44]. While developmentally-inspired bone regeneration approaches such as EBR have shown promise, the difficulty in translating cell-based therapies has led to increased interest in investigating the role of growth factor-enriched MtVs sequestered within these cartilaginous templates for EBR. To the best of our knowledge, this is the first study to bioengineer soft callus-mimetic MtVs through refining the *in vitro* manufacture of MSC-derived cartilage microtissues. Specifically, we demonstrated the tunability of the MtV biofunctionality for EO. Developing bioinspired MtVs provides an “off-the-shelf” nanotherapeutic that can be manufactured from allogeneic cell sources and stored until required clinically. Moreover, due to their nanosize, this broadens the clinical applicability of these nanotherapeutics beyond bone fracture repair such as implant coatings or the treatment of systemic diseases such as OP. The possible integration of EVs with biomaterials presents an exciting opportunity to create advanced functional materials for bone regeneration, allowing for the controlled release of bioactive molecules while the scaffold provides structural support[26]. These findings provide a substantial shift in improving the clinical realization of EBR-inspired approaches for bone tissue regeneration. The importance of TGFβ and BMP signalling in cartilage and bone development has been well reported in the literature[45, 46]. TGFβ is critical for the early differentiation of MSCs into chondrocytes, by upregulating markers such as Sox9, Col2a1 and aggrecan. As chondrocytes mature, TGFβ maintains the stable cartilage matrix and inhibits hypertrophic differentiation. During terminal differentiation and EO, TGFβ promotes chondrocyte apoptosis [47, 48]. On the other hand, BMP signalling enhances chondrocyte proliferation by increasing Indian Hedgehog (IHH) expression and inhibiting FGF signalling[49, 50]. It also promotes chondrocyte hypertrophy by upregulating Runx2 expression [51]. Both BMP and TGFβ signalling pathways converge to promote Sox9 expression or activity, favouring cartilage matrix production[52]. Harnessing these developmentally important cytokines to bioengineer hypertrophic microtissues *in vitro* is paramount for EBR approaches. This bio-inspired strategy and the importance of tuning the cartilage template’s maturation stage have been demonstrated in previous studies. For instance, de Silva *et al*. showed that goat-derived MSC cartilage microtissues cultured with BMP2 and TGFβ exhibited a more hypertrophic chondrocyte phenotype compared to BMP-free cultures[38]. When implanted within a critical-sized goat iliac wing defect, the microtissues cultured with BMP2 stimulated superior bone regeneration. These findings showcase the importance of tuning the hypertrophic phenotype to stimulate effective bone regeneration. In this study, we validated whether the synergistic effects of TGFβ and BMP2 exposure during chondrogenic differentiation of human-derived stem cell microtissues enable tailoring of the hypertrophic phenotype. Our findings showed enhanced expression of cartilage markers (i.e. GAGs, collagen type II) in BMP2-treated microtissues, consistent with previous findings using goat or rat MSCs[38, 53]. This indicates the plasticity of this treatment strategy across various species. Synergistic BMP2 and hypertrophic conditioning resulted in microtissues exhibiting a more pronounced hypertrophic phenotype (i.e. increased expression of VEGF, collagen type I, collagen type X, ALP) compared to BMP2-free and chondrogenic groups. This aligns with previous research and underscores that the switch from chondrogenic to hypertrophic phenotype is associated with elevated ALP activity and collagen type X production[38]. The presence of collagen type I within BMP2/hypertrophic microtissues might be associated with the transdifferentiation of chondrocytes into osteoblast-like cells[54, 55]. Importantly, the expression of hypertrophic chondrogenic markers *in vitro* has been linked to improved EBR capacity[14, 56]. Taken together, our findings showcase the potential for tuning the chondrogenic/hypertrophic phenotype of stem cell-derived cartilage microtissues, crucial for tailoring and optimizing the biological functionality of derived MtVs for bone regeneration.

A growing body of evidence suggests that biological cues embedded within the cartilage ECM are important for driving ossification *in vivo*. Our group has previously demonstrated the superior bone regeneration within a critical-sized femoral defect in rats induced by allogeneic, devitalized callus mimetics when compared to their living counterparts[53]. This suggests that extracellular bioactive factors play a role in stimulating improved regeneration. Indeed, EVs have been reported to play a crucial role in modulating tissue development and homeostasis due to their diverse bioactive factors[41, 57]. The diversity of growth factors in EVs provides a multi-targeted, holistic approach to stimulate effective bone regeneration, which involves biological processes such as immunomodulation, vascularization and osteogenesis[58]. Thus, we initially investigated how refining chondrogenic differentiation conditions influences the generation of MtVs within these callus mimetics. By employing a well-established protocol of ECM digestion and differential ultracentrifugation, we first aimed to confirm the nanoparticles obtained were EVs by following the MISEV2023 guidelines[59]. Our findings showed that the microtissue-derived nanoparticles exhibited a typical spherical morphology, size distribution and tetraspanin positive markers indicative of EVs[60, 61]. We also demonstrated the influence of different chondrogenic differentiation conditions on the quantity of MtVs produced. Through EV protein quantification, NTA and CD63 ELISA, we found that BMP2 conditioning significantly improved the quantity of vesicles embedded within the microtissues when compared to the TGFβ group alone. The BMP2-treated groups exhibited a substantially greater microtissue protein content with no significant differences in cell number (i.e. DNA content). This suggests that increased microtissue ECM proteins sequester a higher quantity of MtVs compared to the BMP2-free microtissues. Moreover, characterization of the EV parental microtissues clearly showed the impact of BMP2 treatment on inducing a more hypertrophic phenotype. The altered composition of the ECM (i.e. increase in collagen type I and type X) within these microtissues may contribute to the differential quantity/bioactivity of MtVs observed between the groups treated with or without BMP2. For instance, Kirsch *et al.* (1994) reported the importance of MtV interaction with collagen type II and X in activating calcium influx within these vesicles when compared to matrix-free MtVs[62], thus indicating the role of ECM proteins on regulating MtV function. The differential binding of MtV-associated growth factors to the cartilaginous ECM likely potentiated the differentiation towards a hypertrophic phenotype by providing a reservoir of phenotype-specific bioactive molecules. Our findings also showed a similar correlation in the quantity of sEVs released into the conditioned medium from these different groups. This indicates that by refining the hypertrophic differentiation conditions, we can augment the production of EVs (MtVs and sEVs) from the cells embedded within the cartilage matrix. These findings clearly demonstrate the impact of hypertrophic conditioning on modulating the production of MtVs embedded within these callus mimetics. Accordingly, the correlation between increased hypertrophic differentiation state and MtV quantity likely contributed to the differential regeneration observed between the TGFβ and TGFβ/BMP2 cartilage microtissues when implanted within a critical-sized bone defect model in goats[38].

One of the major defining characteristics of MtVs is their capacity to induce mineralization of the ECM[44, 63]. In the context of EO, the calcification of the cartilage matrix is critical for effectively remodelling this cartilaginous template into bone tissue[64]. ALP is a key protein involved in the mineral nucleation initiated by MtVs. It is an ecto-enzyme located on the outer leaflet of the membrane that can degrade pyrophosphate (inhibitor of mineralization) into monophosphate ions required for crystal growth[65]. Our findings confirmed that both L-MtV and S-MtV subpopulations derived from all microtissue groups exhibited ALP activity. BMP2 exposure significantly enhanced MtV ALP activity (3.24 - 5.4-fold) compared to the BMP2-free groups. Conditioning in hypertrophic medium improved MtV ALP activity (1.22 - 2.02-fold) compared to vesicles obtain from chondrogenic cultures. These findings reveal the importance of BMP2 exposure compared to hypertrophic medium conditioning in upregulating MtV ALP activity. BMP2 has been reported to induce osteogenesis via upregulating the expression of the transcription factor ATF6, activating downstream targets such as ALP[66]. Additionally, we observed that synergistic BMP2 and hypertrophic microtissue conditioning resulted in a substantial increase in MtV ALP activity compared to BMP2 (1.45 - 2.02-fold) or hypertrophic (3.68 - 5.38-fold) groups alone. The differential MtV ALP activity observed in this study is consistent with the ALP immunostaining of the cartilaginous microtissues, indicating a correlation between the parental tissue phenotype and the bioactivity of their MtVs. These findings also correlate with studies investigating the composition of MtVs obtained from different regions of the growth plate. For instance, MtVs derived from chondrocytes in the pre-hypertrophic/upper hypertrophic zone exhibited increased ALP activity compared to vesicles from chondrocytes in the resting zone[67]. In addition to MtV enrichment in ALP activity, these nanoparticles are known to capture extracellular calcium ions to facilitate calcification of the ECM. This occurs through the active transport of calcium ions into the lumen of vesicles through channelling proteins such as annexins [68] or chelation by lipids[30]. Our findings showed that in a cell-free environment, MtVs were able to sequester extracellular calcium ions, substantially increasing the calcification of the collagen type I matrix, consistent with results in the literature[39]. To investigate whether MtVs directly contribute to ECM mineralization through the delivery of mineral components, we quantified their calcium phosphate content. Our analysis revealed that all MtV groups contained calcium phosphate levels comparable to MtV-free controls. This finding suggests that the enhanced mineralization observed in both the acellular calcification model and MtV-treated hBMSCs is unlikely to result from direct deposition of mineral cargo. Instead, it points toward a mechanism whereby MtVs facilitate calcium influx, potentially via membrane-associated proteins such as Annexins, and deliver osteoinductive molecules, including BMP2 and ALP, to promote osteogenic differentiation in recipient hBMSCs. To further interrogate this, we examined matrix mineralization within the microtissues from which the MtVs were derived. As EV cargo is known to reflect the phenotype and activity of their parent cells or tissues[21], minimal mineral staining in these microtissues reinforces the idea that MtVs are not simply carriers of pre-formed mineral. Rather, their role appears to be the delivery of bioactive cues that stimulate osteogenesis in target cells. Collectively, these findings provide evidence that these hypertrophic MtVs act not as passive mineral carriers, but as active mediators of osteoinduction.

In addition to imbuing the MtVs with mineralization bioactivity, both ALP and Annexin proteins have been reported to facilitate the binding of these nanoparticles to collagen-rich ECMs[68–71]. Thus, the upregulation in collagenous proteins observed within the hypertrophic microtissues (i.e. collagen type I, II and X), may contribute to sequestering MtVs with enhanced mineralization bioactivity. Moreover, besides their role in stimulating ECM mineralization, the delivery of both ALP and Annexin proteins to recipient cells may facilitate lineage-specific differentiation[28]. Taken together, these results showcase that by refining the chondrogenic differentiation conditions, we are able to mimic the distinct maturation states of chondrocytes in the growth plate, resulting in the production of MtVs with enhanced and tunable biomineralization capacity. These ALP-enriched MtVs likely contribute to stimulating recipient cells osteogenic differentiation, ECM binding and mineralization. Moreover, these findings signify the function of the matrix-bound vesicles within the devitalized callus mimetics in facilitating the calcification and remodelling when implanted *in vivo*[38, 53].

Due to their nanosized nature and enrichment with multiple growth factors, the role of EVs in stimulating cellular recruitment has been highlighted as one of their many mechanisms in promoting tissue healing[72, 73]. EVs are naturally recognized by cells and are readily taken up through various endocytic pathways, thus increasing the efficiency of bioactive factor delivery to the target cell[74, 75]. Our findings confirmed that the MtVs derived from all microtissue groups were efficiently taken up by recipient hBMSCs, indicating that the isolation procedure did not damage key proteins involved in the EV uptake machinery. In addition to assessing their internalization by recipient cells, we evaluated the capacity of the MtVs to augment cellular function. The improved proliferation and migration capacity of MtV-treated hBMSCs demonstrates the capacity of the vesicle-associated bioactive factors to stimulate the recruitment of these progenitor cells, an important process in EO. Critically, our results showed enhanced proliferation induced by MtVs derived from BMP2-cultured microtissues. It has been demonstrated that BMP2 can enhance the recruitment of MSCs *in vitro* and *in vivo*. For instance, Liu *et al* (2018) reported that recombinant human BMP2 increased the migration of BMSCs *in vitro* via activating the CDC42/PAK1/LIMK pathway[76]. Moreover, a BMP2-loaded collagen scaffold recruited a higher number of BMSCs injected within the circulatory system in nude mice compared to BMP2-free collagen scaffolds [73]. Therefore, with enhanced BMP2 content within the BMP2/hypertrophic MtVs, quantified via the BMP2 ELISA, this likely contributes to the superior recruitment capacities of these nanoparticles. In the context of EO, the degradation of the hypertrophic cartilaginous matrix *in vivo* likely liberates these entrapped growth factor-enriched vesicles, enabling the recruitment of key cells responsible for tissues remodelling.

In addition to the recruitment of these endogenous progenitor cells, it is important to stimulate their lineage-specific differentiation as effective remodelling of the cartilaginous template into bone tissue is crucial for EBR. Several studies have reported the capacity of EVs to induce the osteogenic differentiation of progenitor cells through the delivery of multiple bioactive factors[41, 61], thus providing a multifactorial holistic approach to stimulate bone regeneration. Therefore, we evaluated the capacity of the cartilaginous microtissue-derived MtVs to stimulate the osteogenesis of recipient hBMSCs. Our results showed that MtVs derived from microtissues of a more mature hypertrophic phenotype (BMP2 and/or hypertrophy) substantially improved hBMSCs osteogenic differentiation and mineralization capacity. These findings confirm the biological potency of the MtVs exhibiting enhanced ALP activity and calcium binding properties. We also showcased that BMP2 conditioning of the microtissues enriched the MtVs in both chondrogenic and hypertrophic groups with higher levels of BMP2 when compared to the vesicles derived from the BMP2-free microtissues, quantified via BMP2 ELISA. Interestingly, there was a greater fold difference in the quantity of BMP2 within the L-MtVs subpopulation (∼23-fold) when compared to the S-MtVs (∼15-fold) derived from BMP2-treated or untreated microtissues. This is probably due to the increased size of the L-MtVs allowing for greater enrichment of protein such as BMP2 when compared to the S-MtVs. Moreover, it is possible that these two MtV subpopulations are generated from different EV biogenesis routes differentially loading BMP2 within these nanovesicles, although this would require further investigation. Although, it is clear that BMP2 conditioning of the parental microtissue enriches the vesicles with enhanced BMP2 content, this does not directly correlate with MtV ALP activity and hBMSCs mineralization observed in this study. It has been reported that MtVs derived from a rat growth plate were enriched with BMP1 through 7, in addition to non-collagenous proteins including VEGF, bone sialoprotein, osteopontin, osteonectin, and osteocalcin[31]. Therefore, this highlights the importance of the diverse biological composition of MtVs in stimulating osteogenesis via a multitargeted holistic approach. Moreover, due to the issues with the controlled release of BMP2 and enhancing the bioactivity of the protein *in vivo*, the BMP2-enriched MtVs in this study may improve the therapeutic efficacy of this growth factor by 1) improving the protein half-life (protection from premature degradation) and 2) delivery of BMP2 directly to cells of interests (eliminating off-target side effects). Importantly, the superior osteoinductive potency of MtVs derived from the hypertrophic microtissues observed in this study was consistent with the enhanced bone formation induced by the BMP-stimulated, devitalized cartilage microtissues observed in a critically-sized iliac wing defect in goats[38].

In the final stages of EO, the avascular cartilage template is replaced by a highly vascularized bone tissue[77]. Hypertrophic chondrocytes are the primary drivers of EO, a process that requires extensive vascularization[78]. Due to the importance of vascularization in EO, hypertrophic chondrocytes likely produce pro-angiogenic factors to facilitate vascular invasion into the cartilage template. It has been reported that MtVs derived from hypertrophic chondrocytes contain pro-angiogenic factors such as VEGF[31]. Moreover, immunohistochemical characterization of the microtissues confirmed enhanced expression of VEGF in the more hypertrophic tissues (BMP2/hypertrophic groups), consistent with previous findings in goat MSC microtissues[38]. Thus, we evaluated whether hypertrophic priming improved the angiogenic capacity of the microtissue-derived MtVs. Our findings showed that EVs isolated from the cartilaginous tissues cultured in BMP2/hypertrophic conditions substantially enhanced angiogenic tube and network formation of hECFCs when compared to the vesicles from either chondrogenic or BMP2-free groups. Moreover, to confirm whether MtVs were differentially enriched with pro-angiogenic growth factors, we quantified the nanoparticles’ VEGF content. Our findings showed that transitioning to a more hypertrophic phenotype through exposure to BMP2 and/or hypertrophic medium, significantly enhanced the enrichment of VEGF within the MtVs when compared to the BMP2- and hypertrophic-free groups. Interestingly, we observed a greater fold change increase in the quantity of VEGF within the MtVs between hypertrophic microtissues with or without BMP2 (∼1.86-fold) when compared to chondrogenic spheroids with or without BMP2 (∼1.25-fold). This highlights the importance of synergistic BMP2 and hypertrophic conditioning on potentiating VEGF expression within the MtVs. This enrichment in MtV-associated VEGF correlated with the enhanced hECFC angiogenic tube formation induced by MtV treatment, although other EV-associated growth factors (i.e. nucleic acids, proteins) likely contributed to stimulating angiogenesis. For instance, BMPs have been reported to promote the angiogenesis of endothelial cells by improving motility and invasion[79–81]. Several studies have reported the superior vascularized bone formation induced by the delivery of multiple growth factors when compared to single growth factor treatments[82, 83]. Thus, the enrichment of both VEGF and BMPs within the vesicles from hypertrophic microtissues, likely synergistically stimulated the enhanced hECFC angiogenesis observed in this study and the improved vascularized bone formation induced by hypertrophic cartilage tissues implanted *in vivo* [38]. The angio-inductive potency of these matrix-bound vesicles signifies their role in stimulating vascularization of the avascular cartilage template during EO. Moreover, this study showcases the tremendous potential of *in vitro* generated MtVs as a tunable nanotherapeutic approach to meet the clinical challenge of reconstructing large-scale bone defects or defects with impaired vascularization.

MtVs have long been recognized as essential mediators of mineralization in developing cartilage and bone [23, 27]. Here, we present a comprehensive proteomic profile of HB MtVs, revealing biological processes, molecular functions, and signalling pathways that highlight their active role in EO. Rather than being passive byproducts of chondrocyte hypertrophy, MtVs are enriched with proteins that drive ECM remodelling, mineral deposition, and cell-matrix communication (Figure 11). GO analysis revealed strong enrichment in processes related to ECM organization, ossification, and bone mineralization, confirming the functional relevance of MtVs in skeletal development and aligning with the stark induction of osteogenic differentiation and mineralization observed in this study. KEGG pathway analysis further reinforces the multifunctional bioactivity of MtVs, revealing significant enrichment in Integrin signalling, Wnt signalling, and PI3K-Akt signalling pathways. These pathways are known to regulate cell survival, proliferation, and differentiation, particularly in the context of chondrocyte maturation and osteogenesis[84–86]. The presence of calcium-binding proteins (ANXA2, ANXA5) and ECM structural components (COL1A1, COL2A1, COL10A1, FN1) aligns with evidence in the literature implicating MtVs as initiators of hydroxyapatite crystal nucleation within cartilage[68, 69]. Interestingly, the detection of components related to mineral absorption and protein digestion pathways also points to a broader functional repertoire, potentially reflecting roles in nutrient regulation or proteolytic processing within the mineralizing front. The overrepresentation of metallopeptidases (MMP2, MMP14) suggests a role for MtVs in localized matrix degradation, potentially facilitating vascular invasion and bone remodelling[27, 87, 88]. The proteomic profile of these vesicles revealed a strong angiogenic signature, underscored by the presence of key regulators of vascular development and remodelling (Figure 11). Angiogenic proteins including VEGFA (a central driver of endothelial proliferation and tube formation), ENG (Endoglin, a TGF-β co-receptor essential for angiogenesis and vascular remodelling), and SPARC (known to modulate cell-matrix interactions and enhance angiogenic signalling) were enriched within the HB MtVs[89–91]. Interestingly, THBS1 (Thrombospondin-1), typically considered an anti-angiogenic factor, was present, suggesting a potential regulatory role in balancing pro- and anti-angiogenic cues. Structural ECM proteins such as FN1 (Fibronectin 1) and COL4A1/COL4A2 (type IV collagen chains) further support endothelial cell adhesion, migration, and the stabilization of nascent vascular structures. Cytoskeletal and membrane-associated proteins including VIM (Vimentin) and Annexins (ANXA2, ANXA5), are implicated in endothelial cell migration, membrane dynamics, and plasmin-mediated matrix remodelling[92–95], reinforce the vesicles’ potential role in promoting vascular invasion and angiogenic remodelling within regenerating tissues. These findings support the notion that MtVs derived from hypertrophic chondrocytes function as bioactive nanostructures that contribute not only to mineral deposition but also to the orchestration of complex signalling events at the cartilage-bone interface.

**Figure 11.**
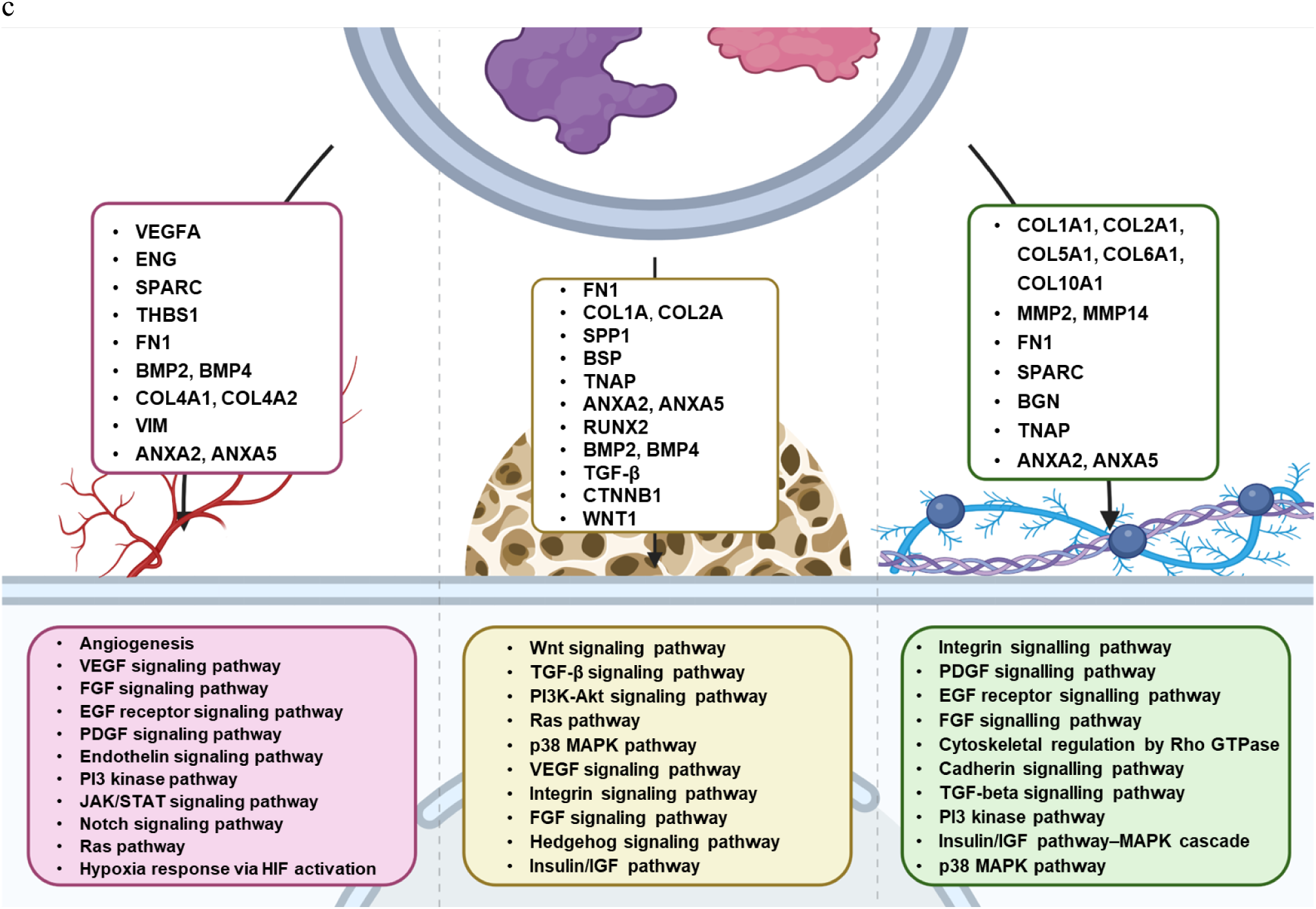
Schematic representation of the multifunctional role of hypertrophic MtVs enriched proteins involved in stimulating angiogenesis, osteogenesis and ECM organization/remodelling.

To further explore the molecular basis of the potent regenerative effects observed *in vitro*, we conducted a protein-protein interaction network analysis using the STRING database, focusing on proteins identified within HB MtV. The resulting network reveals distinct functional clusters with high connectivity, indicating coordinated molecular functions. Notably, a dense core of ribosomal and translational machinery proteins (e.g., RPL, RPS, EIF family proteins) forms a highly interconnected module (highlighted in magenta), suggesting an active role for MtVs in regulating local protein synthesis and potentially facilitating ECM remodelling or cellular translation upon uptake by recipient cells[96–98]. A second major cluster is enriched in ECM-related proteins, including multiple collagens (COL1A1, COL2A1, COL5A1, COL6A1), fibronectin (FN1), SPARC, and BGN, which are central to bone matrix organization[99–102]. The strong connectivity among these ECM proteins supports the structural and signalling contributions of MtVs to osteogenesis. In particular, the presence of COL10A1, a hallmark of hypertrophic chondrocytes, reinforces the developmental origin and EO context of these vesicles. Angiogenic potential is reflected by the inclusion of VEGF pathway-related proteins (e.g., TGFβ1, ENG, SPARC, THBS1) and basement membrane components that facilitate vascular invasion during bone development[103, 104]. Additionally, proteins involved in vesicle trafficking and immune modulation (e.g., ANXA5, VIM, S100A10) appear well-connected, potentially contributing to the MtV-mediated regulation of cellular behaviour in the osteoimmune niche[105, 106]. Notably, mitochondrial and metabolic proteins such as ATP5F1B, PKM, and LDHA may reflect the energy demands of matrix production and mineralization, and their export in MtVs could influence local cellular metabolism or mineral nucleation[107, 108]. *In vitro*, hypertrophic cartilage-derived MtVs markedly enhanced osteogenic differentiation, evidenced by elevated ALP activity, calcium deposition, and upregulation of osteogenic genes. They also promoted endothelial tube formation, aligning with their pro-angiogenic protein cargo. The strong concordance between MtV proteomic profiles and functional outcomes suggests these vesicles are naturally optimized carriers of osteoinductive and angiogenic signals. STRING network analysis further revealed coordinated enrichment in vesicle trafficking, ECM remodelling, ossification, angiogenesis, and metabolic pathways, highlighting the systems-level mechanisms driving their bioactivity.

Due to their nanosized nature and physiochemical stability, these bio-mimetic multifunctional and bio-tunable MtVs could prove beneficial to a wide range of musculoskeletal clinical applications, thus facilitating the realization of EBR-inspired approaches in the clinical arena (Figure 12). The possible integration of designer MtVs into biomaterials represents a significant advancement in the development of smart scaffolds for bone regeneration. Such functionalized materials offer a dual advantage: they provide structural support while actively promoting bone formation through the controlled release of bioactive factors [26, 109]. This controlled release mechanism offers significant advantages over traditional approaches, addressing drawbacks associated with the direct application of osteoinductive growth factors like BMP-2, such as heterotopic ossification[9]. While our study focuses on MtVs, it is important to note the broader context of EV research in bone regeneration. EVs from various sources have shown great potential for biomaterial functionalization. For instance, MSC-derived EVs functionalized to fibronectin-coated decalcified bone matrix scaffolds improved bone regeneration and vascularization in a rat cranial bone defect [110]. Moreover, EV functionalization has shown promise in improving medical devices, for example as coating on titanium implants. Zhai et al. (2020) demonstrated improved osseointegration and bone formation by attaching EVs to polylysine-coated titanium alloy scaffolds via electrostatic interactions[111]. Further, Cheng et al. (2025) reported that functionalization with macrophage-derived EVs enhanced the peri-implant osseointegration of titanium implants under diabetic conditions, highlighting the potential of EV-based strategies to improve implant functionality in disease-specific contexts[112]. Overall, vesicle-based biomaterial functionalization approaches offer a dual promise: they can serve as standalone treatments incorporated into biomaterials and as enhancers of existing medical devices. As standalone therapies, these vesicle-based approaches provide a more targeted and regulated release of growth factors from scaffolds, potentially offering safer and more effective alternatives to traditional bone regeneration methods using for example osteoinductive growth factors. Additionally, by functionalizing titanium screws, spinal cages, and other orthopedic implants with MtVs or EVs, we can enhance their biocompatibility, osseointegration, and overall therapeutic efficacy. This dual role positions vesicle-based approaches at the forefront of innovation in bone regeneration strategies, promising improved outcomes across a wide range of orthopedic applications.

**Figure 12.**
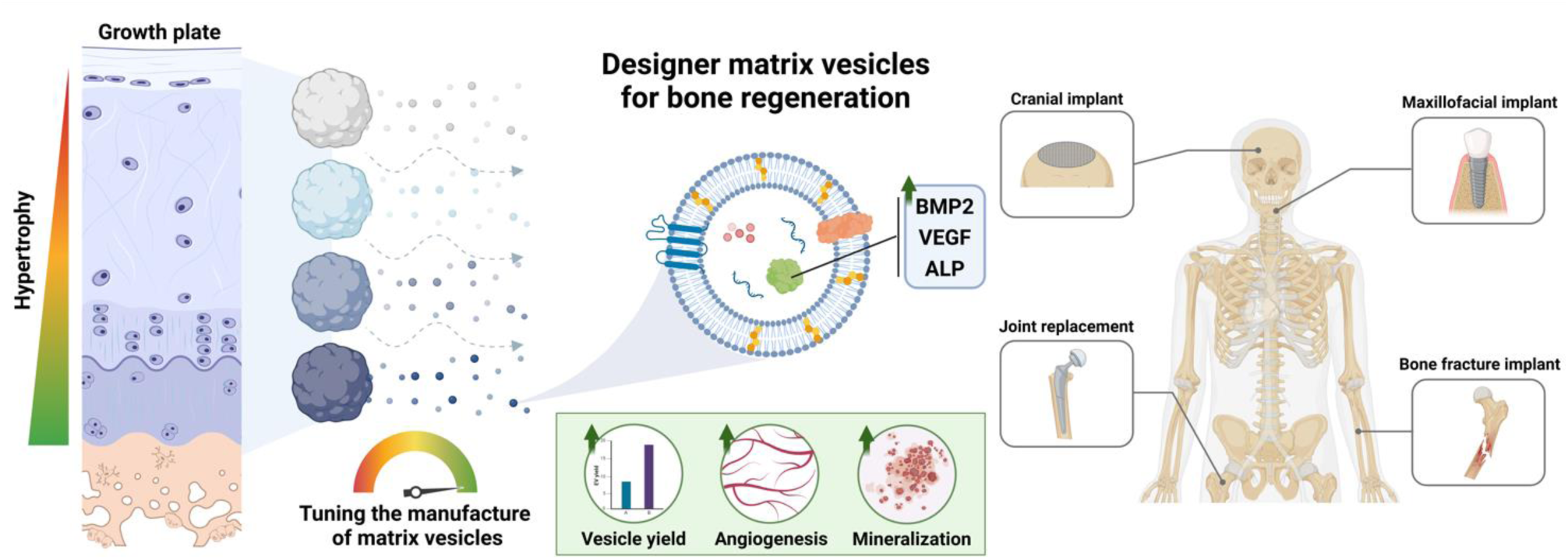
Schematic overview highlighting the potential of harnessing bio-mimetic multifunctional and bio-tunable MtVs for bone augmentation as a standalone therapy or to improve the functionality of existing medical devices.

Taken together, this study demonstrates the importance of harnessing biomimetic cues in generating stem cell-derived hypertrophic cartilaginous microtissues *in vitro*. Moreover, we showcase the feasibly of liberating these matrix-bound growth factor-enriched vesicles for use as an “off-the-shelf” nanotherapeutic that can be manufactured at scale from allogeneic cell sources and stored until required. Additionally, by conducting in depth proteomic analysis of these MtVs, this provides insights into the therapeutic contents and elucidates their subsequent mechanism of action. Although the *in vivo* investigation of MtVs is beyond the scope of this present study, which is focused on determining whether bioengineering hypertrophic cartilage microtissues is a viable bio-mimetic approach for engineering MtVs and elucidating their multi-targeted regenerative function, future studies will investigate the osteoinductive efficacy of these MtVs within an appropriate animal model.

## 5. Conclusion

In summary, we have demonstrated for the first time the impact of tuning the *in vitro* chondrogenic differentiation of stem cell-derived cartilage microtissues towards a hypertrophic phenotype on the production of MtVs with improved therapeutic efficacy for EO. This study highlights the potential role MtVs play in the process of endochondral bone formation during fetal bone development and secondary fracture repair. Thus, these findings showcase the considerable potential of harnessing designer MtVs derived from hypertrophic cartilage microtissues as a nanotherapeutic strategy to promote EBR. As we look to the future, the integration of MtVs into biomaterials and medical devices holds immense promise for advancing EBR-inspired approaches in the clinical arena.

## Acknowledgements

This study was supported by the project XLbone (with project number 19260) of the research program OTP which is (partly) financed by the Dutch Research Council. The antibody against collagen type II (II-II6B3), developed by T.F. Linsenmayer, was obtained from the DSHB developed under the auspices of the NICHD and maintained by The University of Iowa, Department of Biology, Iowa City, IA52242.

## Author contribution

F.S. and K.M. study conceptualization, biological laboratory work and manuscript preparation. P.S.A, R.M.V, H.R.V. proteomics analysis. K.M. supervision. E.B. and D.G. contributed revisions and editing. All authors have read and agreed to the published version of the manuscript.

## Data Availability Statement

The data underlying this article will be shared on reasonable request to the corresponding author.

## Supplementary figures

**Supplementary Figure 1.**
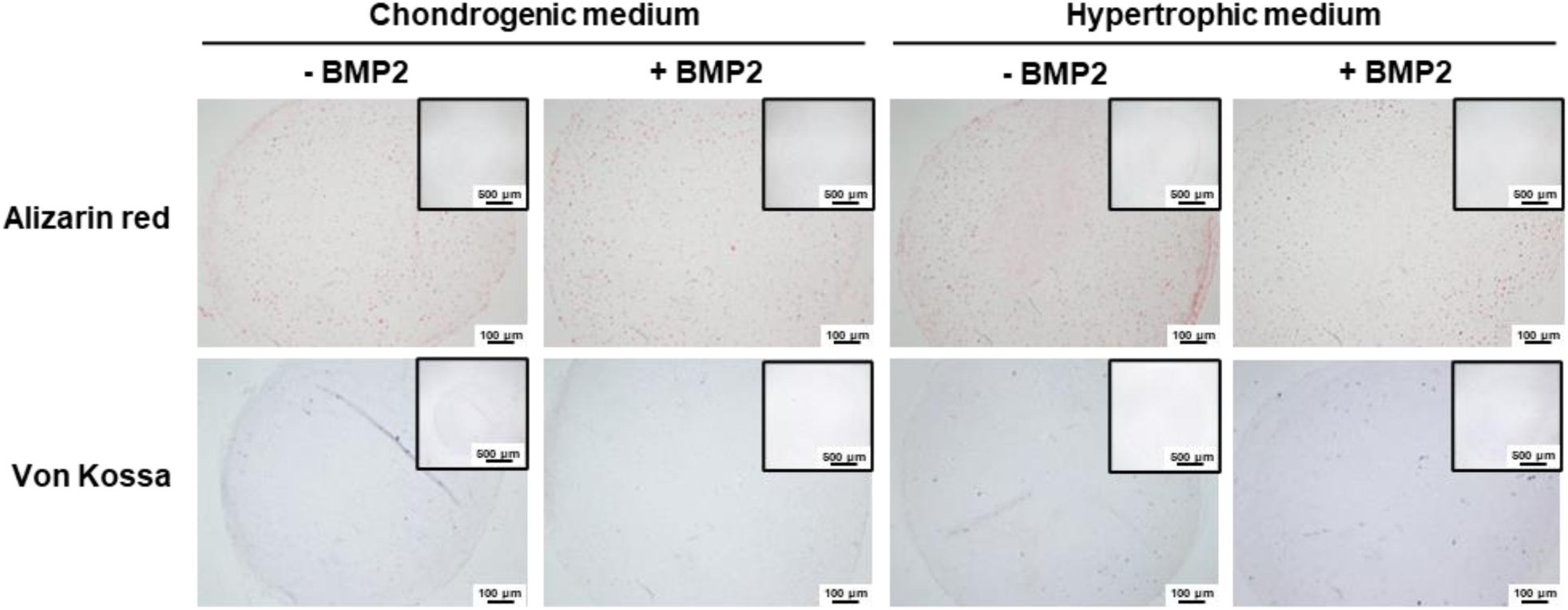
Alizarin red and Von Kossa staining of hBMSC microtissues differentiated with or without BMP2 in chondrogenic or hypertrophic conditions.

**Supplementary Figure 2.**
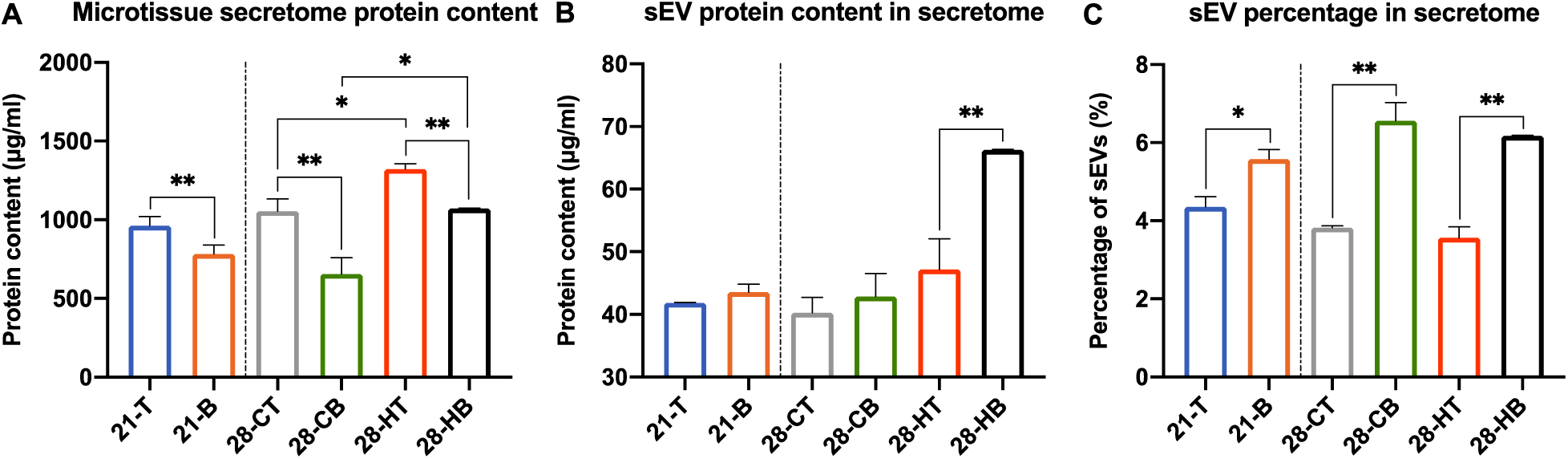
Quantification of A) protein content, B) sEV content, and C) percentage of sEV in the microtissue-secretome at days 21 and 28. Data expressed as mean ± SD (N = 3). *P ≤ 0.05, **P ≤ 0.01 and ***P ≤ 0.001. CT = chondrogenic medium/-BMP2; CB = chondrogenic medium/+BMP2; HT = hypertrophic medium/-BMP2; HB = hypertrophic medium/+BMP2.

**Supplementary Figure 3.**
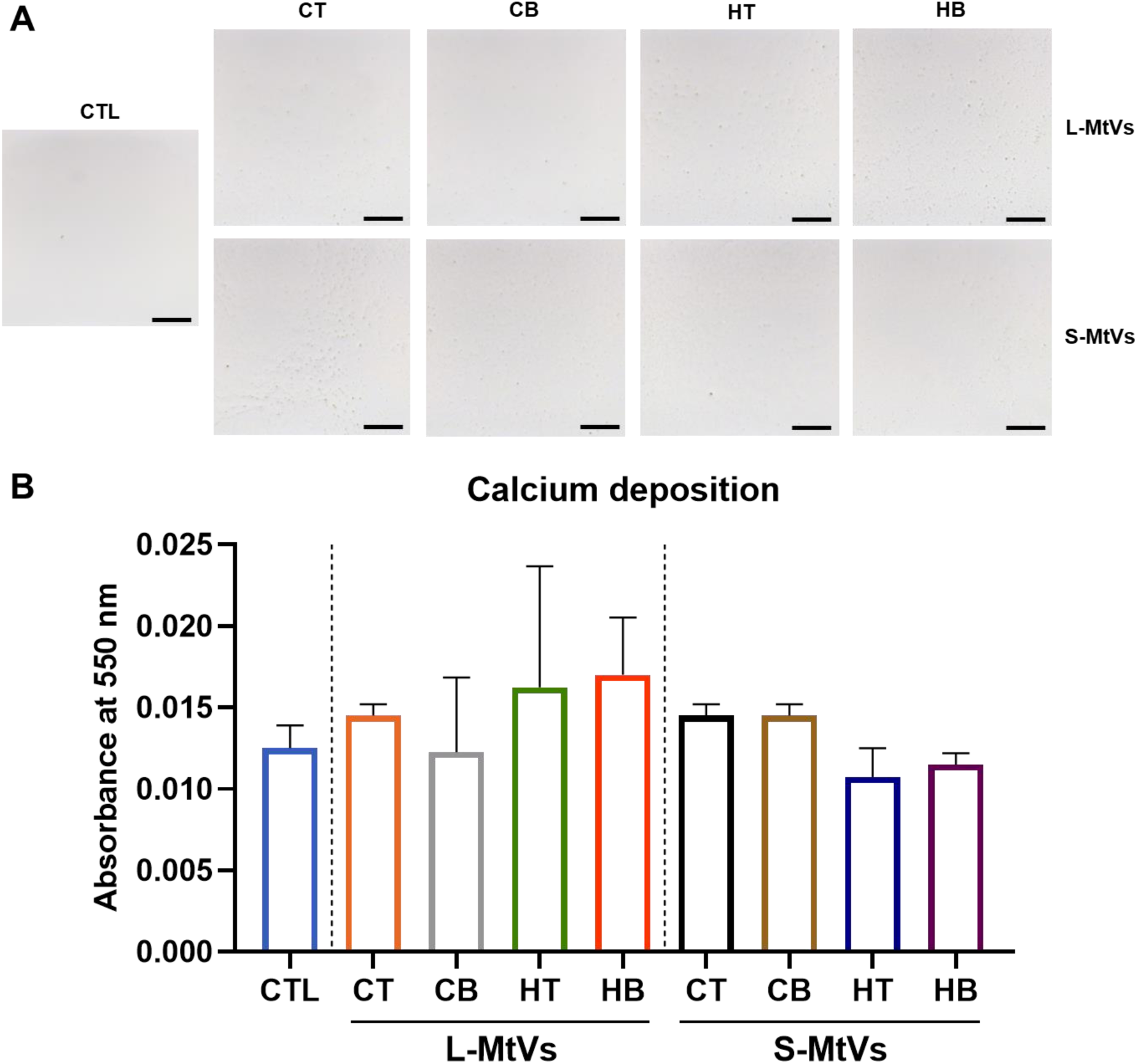
The calcium content of MtVs-coated collagen surfaces. A) Alizarin red S staining for calcium deposition of MtV-functionalized collagen surfaces. Scale bar = 200 µm. B) Semi-quantification of calcium deposition. Data expressed as mean ± SD (N = 3). *P ≤ 0.05, **P ≤ 0.01 and ***P ≤ 0.001. CTL = control; CT = chondrogenic medium/-BMP2; CB = chondrogenic medium/+BMP2; HT = hypertrophic medium/-BMP2; HB = hypertrophic medium/+BMP2.

**Supplementary Figure 4.**
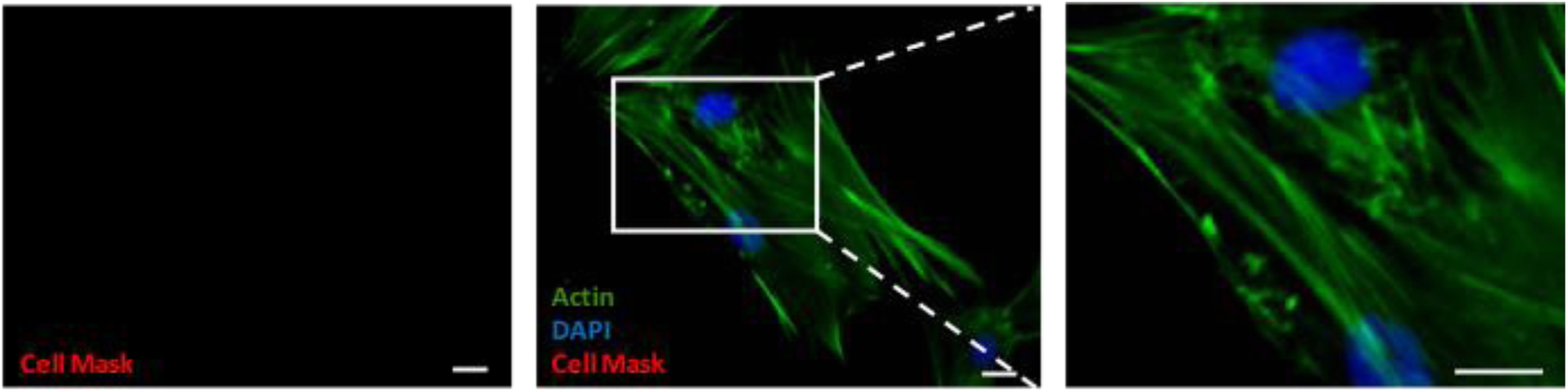
Fluorescence images of non-MtV treated hBMSCs. Scale bar = 10 µm.

**Supplementary Figure 5.**
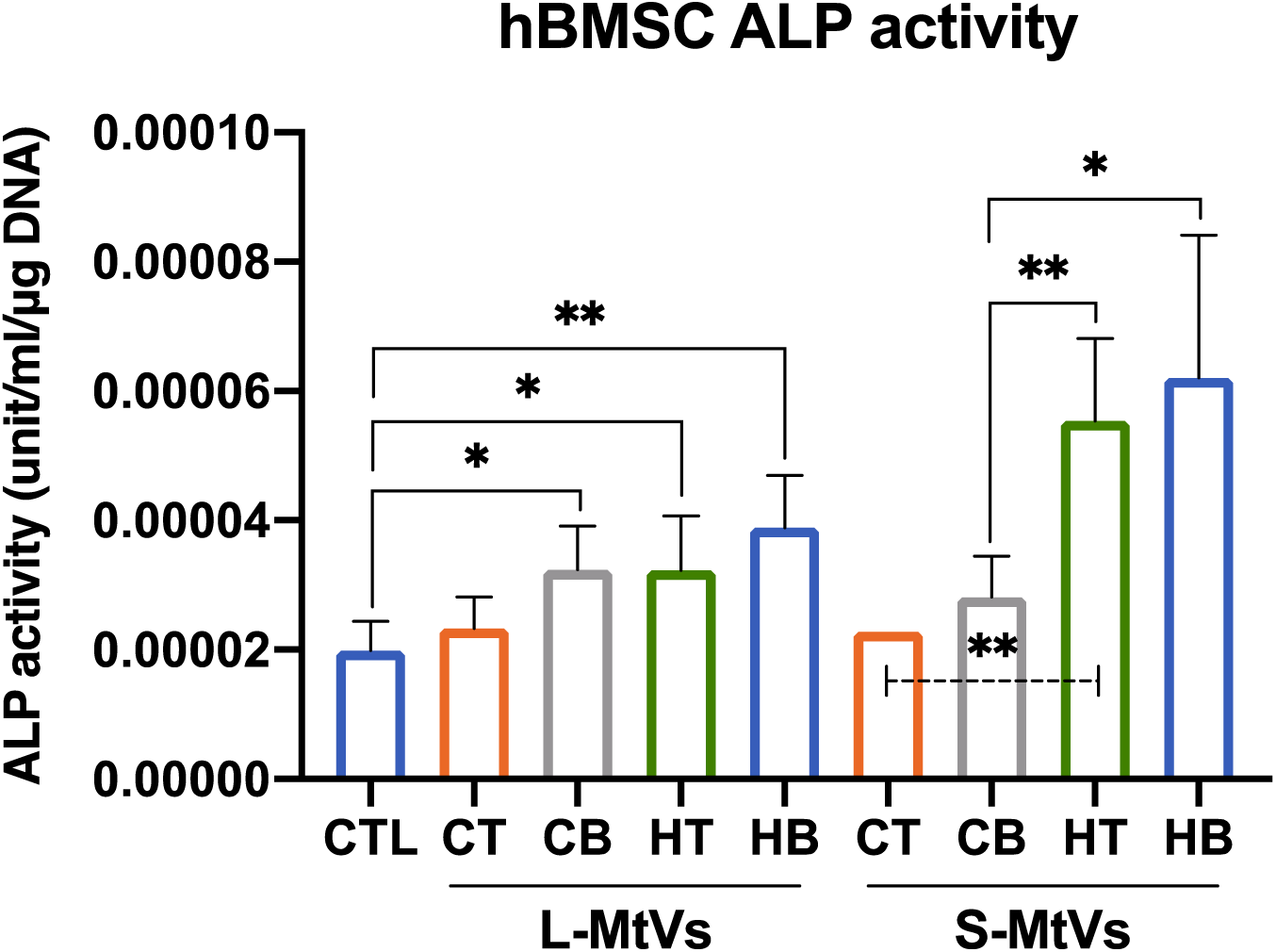
ALP activity of MtV-treated hBMSCs in monolayer following 7 days of osteoinduction. Data expressed as mean ± SD (N= 3). *P ≤ 0.05, and **P ≤ 0.01. CT = chondrogenic medium/-BMP2; CB = chondrogenic medium/+BMP2; HT = hypertrophic medium/-BMP2; HB = hypertrophic medium/+BMP2.

